# Discovery of glycerol phosphate modification on streptococcal rhamnose polysaccharides

**DOI:** 10.1101/337519

**Authors:** Rebecca J. Edgar, Vincent P. van Hensbergen, Alessandro Ruda, Andrew G. Turner, Pan Deng, Yoann Le Breton, Najib M. El-Sayed, Ashton T. Belew, Kevin S. McIver, Alastair G. McEwan, Andrew J. Morris, Gérard Lambeau, Mark J. Walker, Jeffrey S. Rush, Konstantin V. Korotkov, Göran Widmalm, Nina M. van Sorge, Natalia Korotkova

## Abstract

Cell wall glycopolymers on the surface of Gram-positive bacteria are fundamental to bacterial physiology and infection biology. These structures have also gained interest as vaccine antigens, in particular for the human pathogens Group A *Streptococcus* (GAS) and *Streptococcus mutans*. Streptococcal cell wall glycopolymers are considered to be functional homologues of wall teichoic acids but surprisingly lack the biologically-relevant and characteristic anionic charge. Here we identify *gacH*, a gene of unknown function in the GAS Group A Carbohydrate (GAC) biosynthetic cluster, in two independent transposon library screens for its ability to confer resistance to zinc and susceptibility to the bactericidal enzyme human group IIA secreted phospholipase A_2_. To understand the underlying mechanism of these phenotypes, we determined the structure of the extracellular domain of GacH and discover that it represents a new family of glycerol phosphate (GroP) transferases. Importantly, we demonstrate the presence of GroP in both the GAC and the homologous Serotype *c* Carbohydrate (SCC) from *S. mutans,* which is conferred by *gacH* and *sccH* products, respectively. NMR analysis of GAC released from cell wall by non-destructive methods reveals that approximately 30% of the GAC GlcNAc side-chains are modified by GroP at the C6 hydroxyl group. This previously unrecognized structural modification impacts host-pathogen interaction and has implications for vaccine design.

**Graphical abstract:** 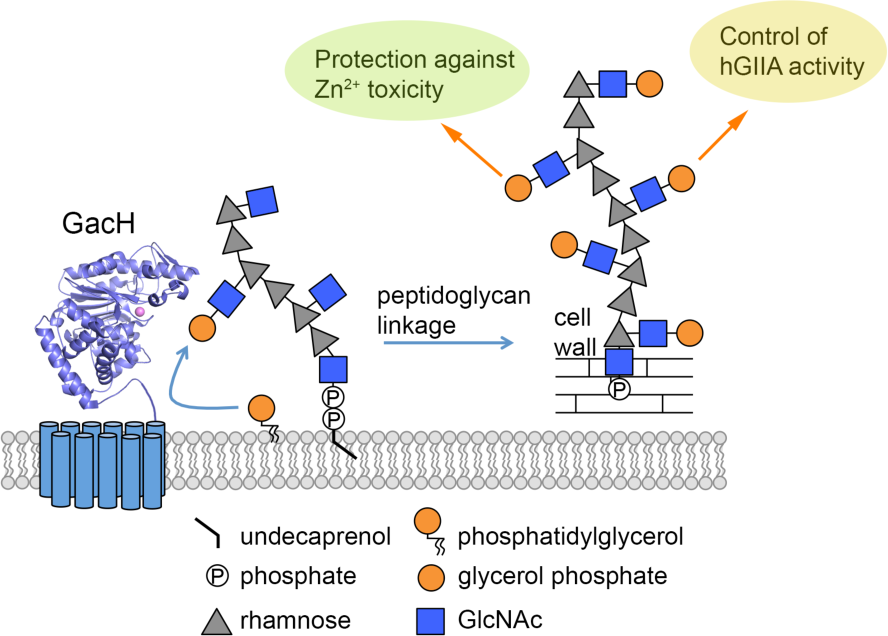

---

Gram-positive bacteria are surrounded by a thick cell wall that consists of a complex network of peptidoglycan with covalently attached proteins and glycopolymers. Cell wall glycopolymers comprise a large family of structurally diverse molecules, including wall teichoic acid (WTA), mycobacterial arabinogalactans and capsular polysaccharides. From these, WTA is perhaps the most widespread and certainly the best-studied molecule. This polyanionic, phosphate-rich glycopolymer is critical for functions such as cell division, antibiotic resistance, metal ion homeostasis, phage-mediated horizontal gene transfer and protection of bacteria from host defense peptides and antimicrobial enzymes ^1–3^. As such, these structures and their biosynthetic pathways are attractive targets for therapeutic intervention, such as antibiotic development and vaccine design. Interestingly, many streptococci lack expression of classical WTA and instead express glycopolymers that are characterized by the presence of L-rhamnose (Rha) ^4^. These structures, referred to as Streptococcal Rhamnose Polysaccharides (SRPs), comprise about 40-60% of the bacterial cell wall mass ^5^, and are historically used for serological grouping of streptococci ^6^. The glycopolymers of two human streptococcal pathogens, Group A *Streptococcus* (GAS; *Streptococcus pyogenes*) and *Streptococcus mutans*, share a characteristic [→3)α-Rha(1→2)α-Rha(1→] polyrhamnose backbone, but are distinguished based on their specific glycosyl side-chain residues, i.e. *N*-acetyl-β-D-glucosamine (GlcNAc) at the C3 position of Rha in GAS ^7^ and α-glucose (Glc) at the C2 position of Rha in *S. mutans c* serotype ^8^. These conserved glycopolymers, referred to as the Lancefield group A Carbohydrate (GAC) and Serotype *c* specific Carbohydrate (SCC), play significant roles in the cell physiology and pathogenesis of GAS and *S. mutans*, respectively. SCC-defective mutants show aberrant cell morphology and division ^9^, increased susceptibility to certain cell wall targeting antibiotics ^10^, defects in biofilm formation ^11^ and reduced ability to induce infective endocarditis compared to the parental strain ^12^. Biosynthesis of the rhamnan-backbone of GAC is essential for GAS viability ^13,14^. Moreover, GAS mutants deficient in the GAC GlcNAc side-chain are viable but more susceptible to innate immune clearance by neutrophils and antimicrobial agents, resulting in significant loss of virulence in animal models of GAS infection ^13,15^. Importantly for both pathogens, GAC and SCC have been evaluated as vaccine antigens. Indeed, immunization with GAC or SCC induces opsonophagocytic antibodies that enhance killing of GAS and *S. mutans*, respectively ^8,16,17^. In addition, GAC has proven efficacious as a vaccine antigen through active immunization in mice ^16,17^.

The GAC and SCC biosynthetic pathways are encoded by 12-gene clusters ^4,13^, herein designated as *gacABCDEFGHIJKL* and *sccABCDEFGHMNPQ* (Fig. 1a), respectively. The first seven genes in both operons are conserved in many streptococcal species and they participate in polyrhamnose backbone synthesis and transport ^18^. In GAS, we have demonstrated that *gacI, gacJ, gacK* and *gacL* encode the machinery to generate and add the immunodominant GlcNAc side-chain to the polyrhamnose backbone ^13,19^. In *S. mutans, sccM* and *sccN* are required for immunoreactivity with serotype *c*-specific antiserum suggesting a function for these genes in Glc side-chain attachment to the polyrhamnose backbone ^20^. In addition to these streptococcal species, similar gene clusters are present in a wide variety of streptococcal, lactococcal and enterococcal species, although the corresponding glycopolymer structures have not all been elucidated ^4^.

**Fig. 1.**
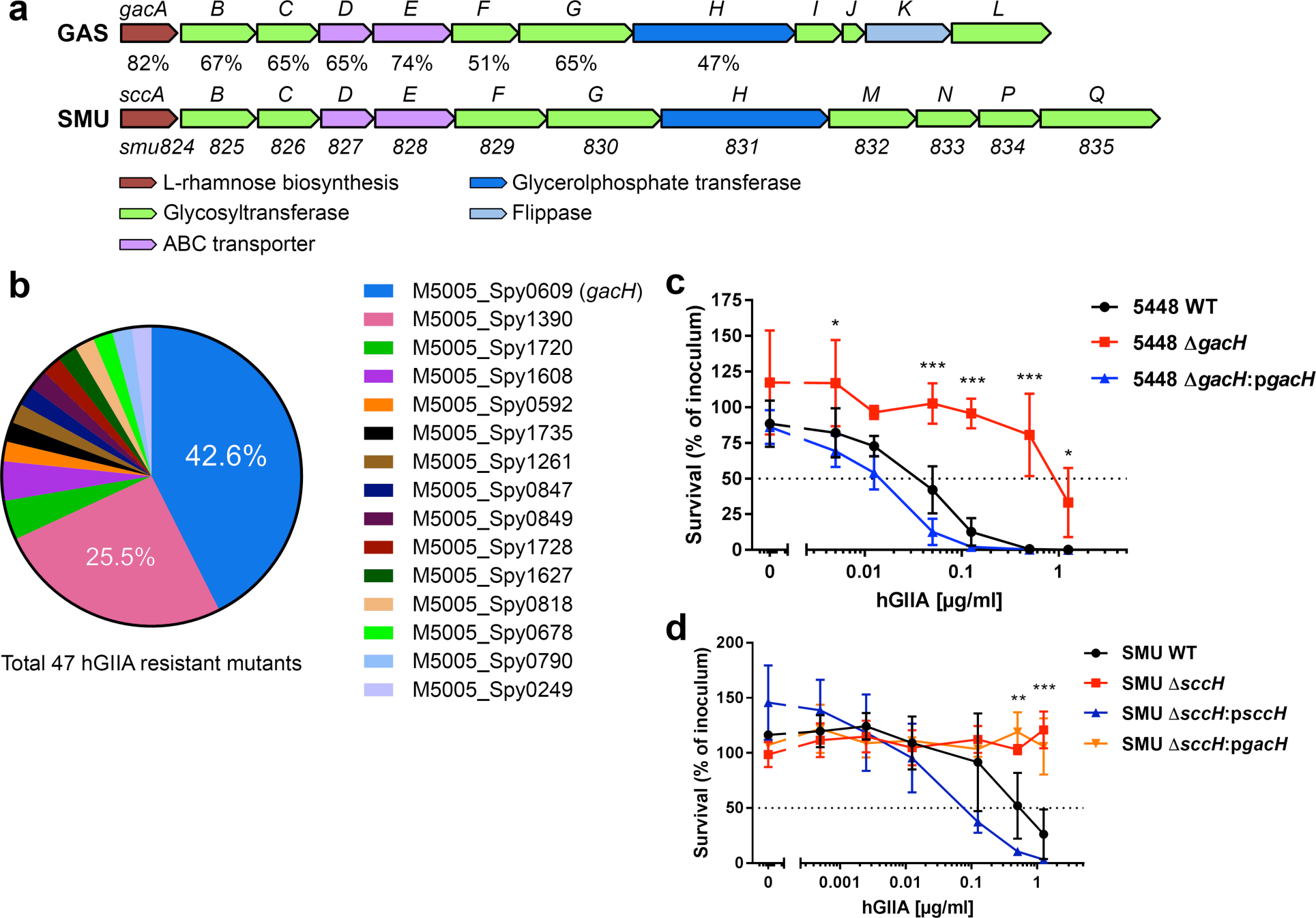
GacH homologues are required for full hGIIA bactericidal activity against GAS and *S. mutans*. (**a**) Schematic representation of GAC and SCC biosynthetic gene clusters. SCC gene cluster *smu*824-835 was renamed *sccABCDEFGHMNPQ*. Sequence identity (%) between homologous proteins is indicated. Sequences of GAS 5005 and *S. mutans* UA159 were used for comparison. (**b**–**d**) *gacH* is identified in Tn-seq screen for hGIIA resistance and its deletion confers resistance to hGIIA. (**b**) Transposon gene location in 47 hGIIA resistant mutants after exposure of *Krmit* mutant transposon library to lethal concentrations of hGIIA. (**c**) Deletion of *gacH* in GAS 5448 and (**d**) the *gacH*-homologous gene *sccH* in *S. mutans* increases hGIIA resistance more than 10-fold. Data represent mean ± standard deviation of three independent experiments. *, *p* < 0.05; **, *p* < 0.01; ***, *p* < 0.001.

Within this wider collection of species, one of the conserved genes present in the SRP biosynthetic clusters, a *gacH* homologue, encodes a putative glycerol phosphate (GroP) transferase of unknown function. Recently, we employed the *Krmit* GAS transposon mutant library ^14^ and identified *gacI* and *gacH* as genes that confer sensitivity of GAS to human group IIA secreted phospholipase A2 (hGIIA) (http://creativecommons.org/licenses/by-nc-nd/4.0/ ^21^, an important bactericidal protein of the innate immune defense against Gram-positive pathogens ^22^. Complementary, we now present the data that *gacH* was the only valid hit when exposing the *Krmit* library to a lethal concentration of hGIIA. Using the same transposon library, *gacH* was also identified as a gene providing resistance to zinc toxicity. In pursuit of the underlying mechanisms for GacH-mediated hGIIA resistance and protection against zinc toxicity, we have characterized the function of GacH at the genetic, biochemical and structural level. Our study identified a previously overlooked GroP modification on both GAC and SCC, and demonstrated a function of GacH homologues in the transfer of GroP to SRP. GroP is attached to the GlcNAc side-chains of GAC at the C6 hydroxyl group as shown by NMR analysis. These new insights into the structure, biosynthesis and function of GAC and SCC justify a re-examination of the chemical structures of cell wall glycopolymers in other streptococcal species. Furthermore, the discovery of GroP modification of GAC and SCC offers the potential for future vaccine development and provides a framework for investigation of the function of this modification in bacteria.

## Results

GacH homologues are required for full hGIIA bactericidal activity against GAS and

*S. mutans*.

We previously identified *gacH* in a GAS transposon library screen against the bactericidal activity of hGIIA ^21^. Complementary to this susceptibility screen, we also exposed the *Krmit* GAS transposon library ^14^ to a lethal concentration of recombinant hGIIA to identify resistant mutants. Only 47 colonies were recovered after exposure. Sequencing identified that 43% of the recovered mutants (20 out of 47) had a transposon insertion in *gacH*, and 26% in *M5005_Spy_1390* (12 out of 47) (Fig. 1b). *M5005_Spy_1390* was also identified in the initial susceptibility screen as an artifact due to biased transposon insertions ^21^ and therefore not followed up further. To validate our finding for *gacH*, we generated a *gacH* deletion mutant in a GAS clinical isolate of the globally-disseminated serotype M1T1 clone 5448, creating 5448Δ*gacH*, and complemented the mutant with *gacH* on an expression plasmid, creating 5448Δ*gacH:*p*gacH*. Exposure of these strains to a concentration range of hGIIA revealed that deletion of *gacH* increased GAS resistance to hGIIA over 10-fold, which was restored back to wild-type (WT) in 5448Δ*gacH:*p*gacH* (Fig. 1c). The *gacH*-mediated hGIIA resistance was also observed in two different GAS backgrounds, 2221 (M1T1 GAS clone strain) and 5005 (clinical *covS* mutant isolate of M1T1 strain) (Supplementary Fig. 1), demonstrating that the effect is conserved across GAS strains of the M1T1 background and independent of CovRS status – a two-component system which regulates about 15% of the genes in GAS ^23^.

Since other streptococcal species possess a genetic homologue of *gacH*, we investigated whether the GacH-dependent hGIIA-resistance phenotype was conserved in different streptococci. To this end, we generated a deletion mutant of the *gacH* homologue *sccH* (previously known as *orf7* ^20^) in serotype *c S. mutans* (SMU) strain Xc, creating SMUΔ*sccH*. Deletion of *sccH* rendered *S. mutans* completely unsusceptible to the tested hGIIA concentrations (Fig. 1d), which was restored to WT level by expression of *sccH* on a plasmid. However, heterologous expression of *gacH* in SMUΔ*sccH* did not restore the hGIIA resistance phenotype, suggesting that the enzymes might target different substrates. Taken together, our data indicate that deletion of *gacH* homologues renders streptococci more resistant to the bactericidal activity of hGIIA and that GacH function is species-specific. Interestingly, lack of the GAC GlcNAc side-chain, by deletion of *gacI*, also increases hGIIA resistance, which is attributed to reduced cell wall penetration of hGIIA ^21^. These data suggest that similar to *gacI*, *gacH* might participate in a tailoring modification of GAC.

### GacH and SccH provide protection from zinc toxicity

Recent evidence indicates that neutrophils deploy zinc poisoning as an antimicrobial strategy against GAS during phagocytosis ^24^. To resist Zn^2+^ toxicity, GAS expresses the zinc efflux system encoded by *czcD* ^24^. To search for additional GAS genes that confer resistance to zinc poisoning, we performed a Tn-seq screen of the GAS *Krmit* transposon library ^14^ using two Zn^2+^ concentrations, 10 and 20 µM, selected based on growth inhibition analysis (Supplementary Fig. 2a). Genomic DNA for Tn-seq analysis was collected after T2 and T3 passages (Supplementary Fig. 2b). In addition to the expected importance of *czcD*, we also observed that *gacI* and *gacH* transposon insertions were significantly reduced in the library (P-value of <0.05) after growth with 20 µM Zn^2+^ in both T2 and T3 passages compared to untreated controls indicating that the genes provide resistance against Zn^2+^ toxicity (Fig. 2a-d).

**Fig. 2.**
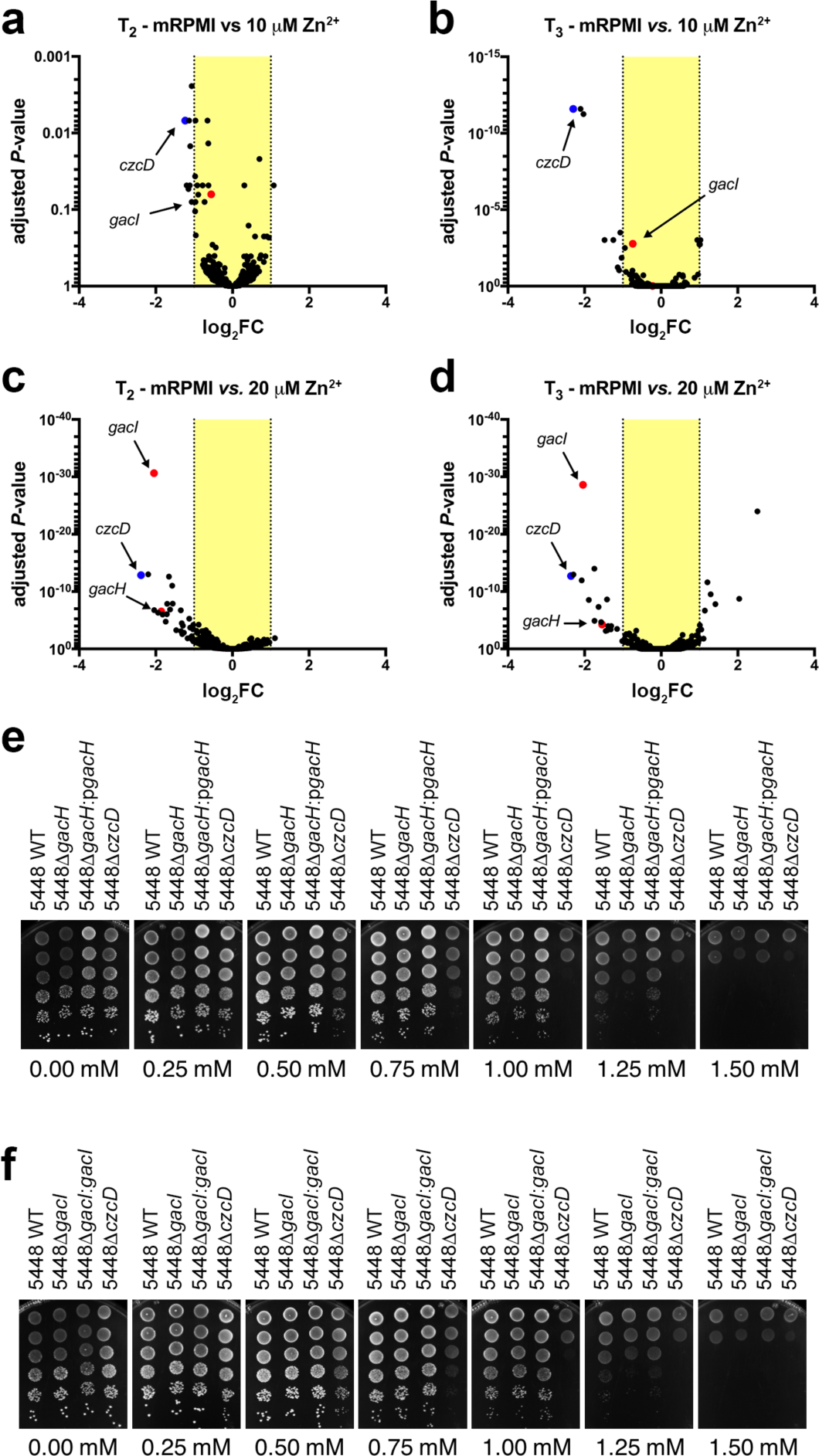
Deletion of *gacI* and *gacH* renders GAS susceptible to Zn^2+^. (**a**–**d**) Tn-seq volcano plots showing representation of *czcD*, *gacH* and *gacI* in GAS *Krmit* transposon library screens for Zn^2+^ tolerance. Log_2_ fold-change (log_2_ FC) in fitness was plotted against adjusted *P*-value from Tn-seq analysis. The outline of the experiment is shown in Supplemental Fig. 2b. Tn-seq screens of the transposon library were conducted using (**a**) 10 µM Zn^2+^ at T_2_, (**b**) 10 µM Zn^2+^ at T_3_, (**c**) 20 µM Zn^2+^ at T_2_, (**d**) 20 µM Zn^2+^ at T_3_. (**e** and **f**) Zn^2+^ sensitivity as tested in drop test assay using strains (**e**) 5448 WT, 5448Δ*gacH* and 5448Δ*gacH:*p*gacH*; and (**f**) 5448 WT, 5448Δ*gacI* and 5448Δ*gacI:gacI*. 5448Δ*czcD* was included as a positive control in both panels. Strains were grown in THY to mid-exponential phase, adjusted to OD_600_ = 0.6, serial diluted and 5 µL spotted onto THY agar plates containing varying concentrations of Zn^2+^. Each drop test assay experiment was performed at least three times.

To confirm that *gacI* and *gacH* are required for GAS resistance to Zn^2+^, we grew 5448Δ*gacH* and 5448Δ*gacI* ^13^ on THY agar supplied with different concentrations of Zn^2+^ (Fig. 2e and 2f). The growth of both mutants was reduced in THY supplied with 1.25 mM Zn^2+^. Expression of full-length *gacH* in 5448∆*gacH,* and *gacI* in 5448∆*gacI* fully complemented the growth phenotypes of the mutants. To investigate whether this was a conserved function for *gacH* homologues, we extended our experiments to *S. mutans*. Similar to the GAS *gacH* deletion mutant, SMUΔ*sccH* was more sensitive to Zn^2+^, in comparison to the parental strain and the phenotype can only be restored by *sccH* but not *gacH* (Supplementary Fig. 3). Hence, our results provide strong evidence that the unknown functions of GacH and SccH are important for protection of streptococci from Zn^2+^ toxicity.

**Fig. 3.**
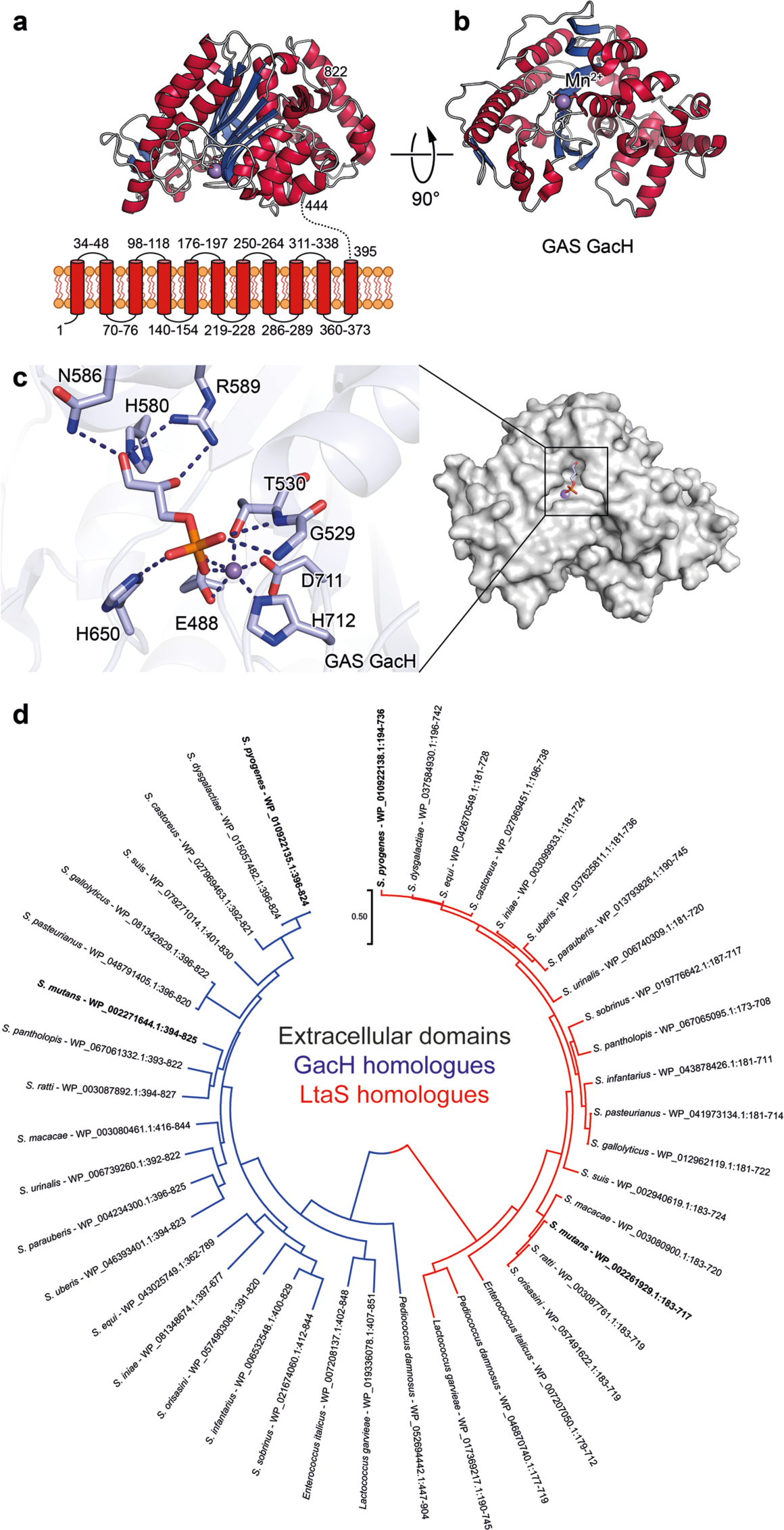
Structure of GacH and phylogenetic analysis of the GacH family of proteins. (**a**) Predicted topology of GacH showing eleven transmembrane helices and structure of extracellular domain with the enzymatic active site oriented towards the cell membrane. (**b**) Structure of apo eGacH viewing at the active site with the Mn^2+^ ion shown as a violet sphere. (**c**) A close-up view of the active site GacH crystal structure in complex with *sn*-Gro-1-P. (**d**) Phylogenetic analysis by Maximum Likelihood method of the predicted extracellular domains of GacH and LtaS enzymes. A phylogenetic tree was generated using the predicted extracellular domains of 21 GacH (blue) and 21 LtaS (red) homologues from the same species. The GAS and *S. mutans* proteins analyzed in this study are indicated in bold. The phylogenetic tree is drawn to scale as indicated by the scale bar, with branch lengths measured in the number of substitutions per site.

### GacH structure

GacH is predicted to contain eleven transmembrane segments in its N-terminal domain, and a C-terminal extracellular domain (eGacH), that is likely to perform the enzymatic function. To gain insight into GacH function, the extracellular domain of GacH (eGacH) was expressed and purified from *E. coli* and its crystal structure was determined in apo form (PDB ID 5U9Z) at 2.0 Å resolution (Fig. 3a). Additionally, to test the hypothesis that GacH is a GroP transferase, we solved the structure of eGacH in complex with GroP (PDB ID 6DGM) at 1.49 Å resolution (Fig. 3c). The apo- and GroP-containing eGacH structures belong to different crystal forms, with two molecules in the asymmetric unit. Analysis of the dimer interface and other crystal contacts revealed that the dimer interface has the largest surface of all the crystal contacts (1809 and 1894 Å^2^ in the two crystal forms). However, it is scored below the stable complex formation criteria, and recombinant eGacH behaves as a monomer in solution. This does not exclude a possibility of a dimer formation in the context of the full-length GacH. The structures of the apo- and GroP-bound eGacH monomers are very similar with root mean square deviation of 0.3 Å for 380 superimposed Cα atoms, as well as between the non-crystallographic copies.

eGacH has an α/β core structure that is similar of the sulfatase protein family, with the closest similarity to lipoteichoic acid (LTA) synthase LtaS ^25,26^ and LTA primase LtaP ^27^ (Supplementary Table 1). LtaS and LtaP are GroP transferases that participate in biosynthesis of LTA, a crucial cell envelope constituent of Gram-positive bacteria. LTA is an anionic polymer consisting of a polyglycerol-phosphate backbone linked to a glycolipid membrane anchor ^28^. The catalytic site of GacH contained a Mn^2+^ ion coordinated by residues E488, T530, D711 and H712, equivalent to residues E255, T300, D475 and H476 of *Staphylococcus aureus* LtaS (Fig. 3c, Supplementary Fig. 4d and 5). The structure of GacH in complex with GroP revealed the position of the ligand in the active site with the phosphoryl group oriented towards the Mn^2+^ ion, and coordinated by residues G529, T530 and H650 (Fig. 3c). The glycerol 2- and 3-hydroxyl groups form hydrogen bonds with side-chains of residues R589, H580 and N586. The positions of GroP and coordinating residues are similar in eGacH and *S. aureus* LtaS structures. For example, the glycerol moiety forms hydrogen bonds with residues H580 and R589 in GacH and equivalent residues H347 and R356 in *S. aureus* LtaS (Fig. 3c and Supplementary Fig. 4d) ^25^. Thus, the structure of GacH in complex with GroP is consistent with the idea that GacH and LtaS use related catalytic mechanisms of GroP transfer to substrates.

**Fig. 4.**
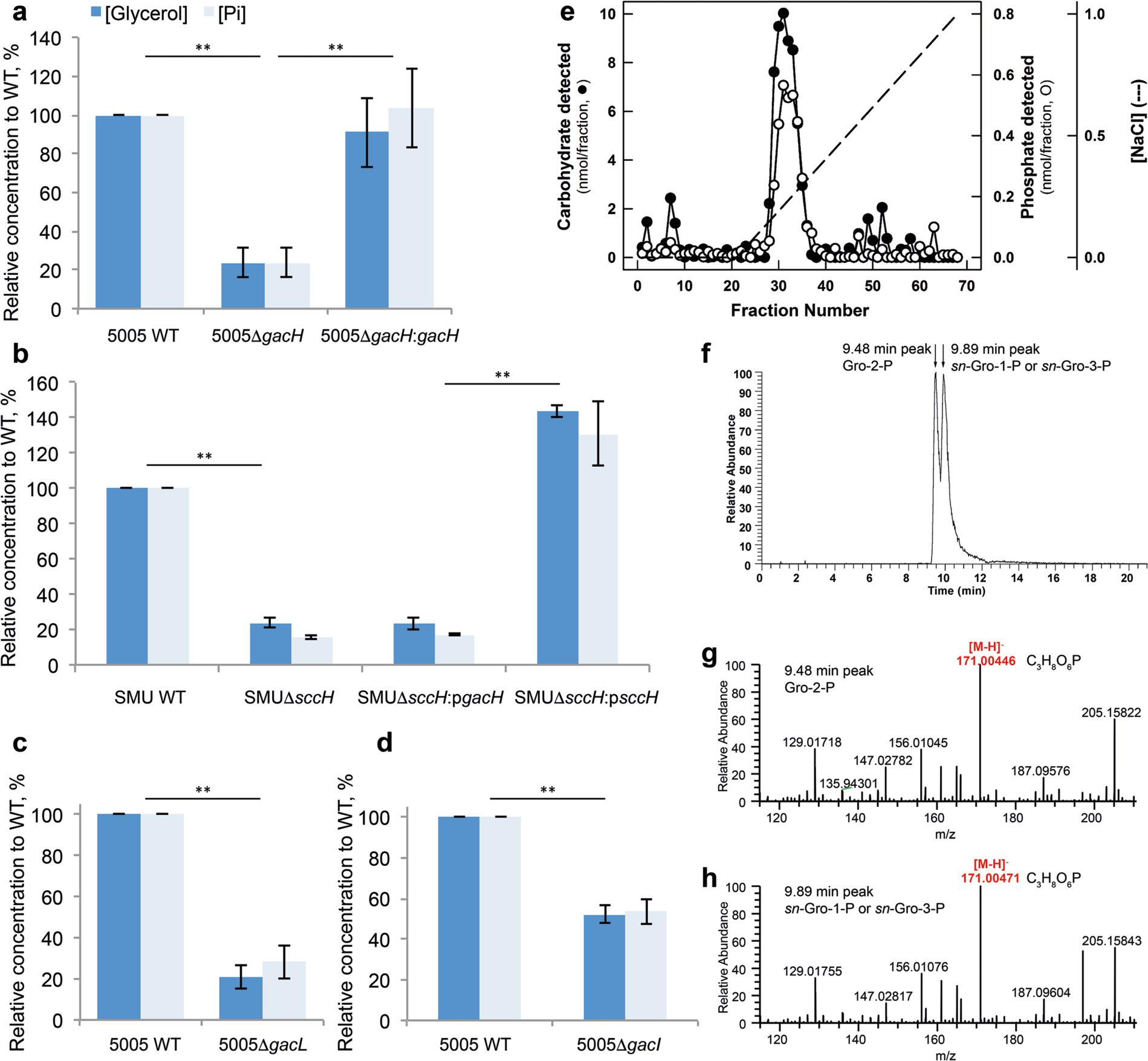
GacH and SccH modify SRPs with *sn*-Gro-1-P. (**a**-**d**) Analysis of glycerol and phosphate content in GAC and SCC isolated from (**a**) GAS 5005 WT, Δ*gacH* and complemented strain; (**b**) *S. mutans* WT, Δ*sccH*, and Δ*sccH* complemented with *sccH* or *gacH*, and (**c** and **d**) GAS 5005 mutants Δ*gacL* and Δ*gacI* that are devoid of GlcNAc side-chain. GAC and SCC were released from bacterial cell wall materials by PlyC and mutanolysin digestion, respectively, and subjected to acid hydrolysis as described in Methods. Phosphate was released from these samples by digestion with alkaline phosphatase and measured using the malachite green assay. Glycerol was measured using a colorimetric glycerol assay kit. The concentration of phosphate and glycerol is presented relative to the WT strain. Data are mean ± standard deviation of three independent replicates. **, *p* < 0.01. *P*-values are shown for glycerol and phosphate concentrations. (**e**) DEAE-Sephacel elution profile of GAC. Isolated GAC was loaded onto an 18 mL column of DEAE-Sephacel. The column was eluted with a 100 ml gradient of NaCl (0-1 M). Fractions were analyzed for carbohydrate by anthrone assay **(•)** and phosphate by malachite green assay (O). (**f**–**h**) Identification of the enantiomeric form of GroP associated with GAC. (**f**) The GroP isomers were recovered from GAC following alkaline hydrolysis and separated by liquid chromatography as outlined in Methods. The elution positions corresponding to standard Gro-2-P and *sn*-Gro-1-P/*sn*-Gro-3-P are indicated by the arrows. LC-MS analysis identifies two extracted ion chromatogram peaks for the molecular GroP ion *m/z* 171.004 [M-H]^-^, which eluted at (**g**) 9.48 and (**h**) 9.89 min. Based on the accurate mass and retention times, these two peaks were assigned as Gro-2-P and *sn*-Gro-1-P/*sn*-Gro-3-P respectively by comparison with authentic chemical standards.

### GacH homologues form a distinct clade of GroP transferases

Taking into consideration GacH structural homology to LtaS, we examined the distribution of the GacH family of proteins throughout the bacterial kingdom and compared the evolutionary relatedness of GacH with LtaS. Interestingly, with a few exceptions, GacH homologues were found predominantly in streptococcal species (Fig. 3d, Supplementary Fig. 4e). In contrast to GacH that contains eleven predicted transmembrane segments, LtaS is composed of an N-terminal domain with five transmembrane helices ^25-27,29^. Based on just the extracellular domains of the proteins, GacH and LtaS-related proteins are grouped in separate clades on a phylogenetic tree, suggesting that the proteins may fulfill distinct functions in bacteria by transferring GroP to different substrates (Fig. 3d).

### GacH homologues decorate respective SRPs with GroP

The genetic, bioinformatic and structural evidence presented in the preceding sections strongly suggest that *gacH* and *sccH* encode novel GroP transferases of unknown function in GAS and *S. mutans* (Supplementary Fig. 6). The presence of these genes in the GAC and SCC biosynthetic clusters implies that they may participate in polysaccharide synthesis in a previously undefined manner. To investigate the possibility that GacH and SccH function in the modification of the respective SRPs with GroP, we enzymatically released SRP from purified cell walls (free of LTA, lipids, proteins and nucleic acids) from GAS 5005, *S. mutans* WT, and corresponding *gacH* and *sccH* deletion strains by treatment with peptidoglycan hydrolases (as described in Methods). Subsequently, the enriched polysaccharide preparations were analyzed for glycerol and phosphate. Hydrolysis with HCl (2 N HCl, 100 °C, 1 hr) released a significant amount of glycerol from GAC and SCC isolated from WT bacterial strains (Fig. 4a, b, and Supplementary Fig. 7). Furthermore, we detected high levels of inorganic phosphate after incubation of these acid-treated samples with alkaline phosphatase (Fig. 4a, b). Of note, the treatment of intact GAC and SCC with alkaline phosphatase alone did not release detectable levels of phosphate (Supplementary Fig. 7), indicating that the phosphoryl moiety is present as a phosphodiester, presumably as GroP. In contrast to WT GAC and SCC, the SRPs isolated from the *gacH* and *sccH* mutants (5005Δ*gacH* and SMUΔ*sccH*, respectively) contained a significantly reduced amount of GroP (Fig. 4a, b). Genomic complementation of 5005Δ*gacH* phenotype by expression of *gacH* on the mutant chromosome restored the WT levels of GroP in GAC (Fig. 4a). Similarly, complementation of SMUΔ*sccH* with plasmid-expressed *sccH* restored GroP incorporation of the mutant to the level of the parental strain (Fig. 4b). In contrast but in accordance with our functional data, expression of *gacH* did not restore the GroP levels in SCC of SMUΔ*sccH* (Fig. 4b). Importantly, analysis of the glycosyl composition of cell walls purified from the GAS and *S. mutans* strains demonstrated that the absence of GacH and SccH did not affect the Rha/GlcNAc and Rha/Glc ratios, respectively (Supplementary Fig. 8). Since the differences in GroP content for 5005Δ*gacH* and SMUΔ*sccH* were not due to changes in the composition of GAC and SCC, our results are consistent with a role for SccH and GacH in modification of SRPs by GroP.

Both structurally and functionally, we can only complement the SMUΔ*sccH* mutant by expression of the native *sccH*, suggesting that the site of GroP attachment to SRPs might involve the species-specific side-chains (Glc vs. GlcNAc), rather than the identical polyrhamnose backbone. Consistent with this hypothesis, the glycerol and phosphate contents in the GAC isolated from two GlcNAc-deficient mutants, 5005Δ*gacL* and 5005Δ*gacI* ^19^ were significantly reduced, similarly to 5005Δ*gacH* (Fig. 4c, d).

To prepare bacterial polysaccharide for more detailed analysis, GAC was released from isolated cell walls by peptidoglycan hydrolase treatment and partially purified by a combination of size exclusion chromatography (SEC) and ion-exchange chromatography (Fig. 4e, Supplementary Fig. 9a). Fractions from both chromatography analyses were assayed for Rha and total phosphate. The majority of the rhamnose- and phosphate-containing material was bound to the ion-exchange column and eluted as a single coincident peak (Fig. 4e). Similarly prepared GAC purified from 5005Δ*gacH* did not bind to the column (Supplementary Fig. 9b). Interestingly, the GAC from 5005Δ*gacH* does appear to contain a small amount of phosphate, although its phosphate content is much lower than the GAC isolated from the WT strain. This data directly supports the conclusion that GAC is modified with GroP donated by GacH.

### Identification of the enantiomeric form of GroP associated with GAC

LTA is formed by the sequential addition of *sn*-Gro-1-P groups transferred by LtaS from the head group of the membrane lipid phosphatidylglycerol ^30,31^. In contrast, the *Bacillus subtilis* poly-GroP backbone of WTA consists of *sn*-Gro-3-P repeats that are synthesized on the cytoplasmic surface of the plasma membrane from CDP-glycerol before export to the periplasm and attachment to the cell wall ^1^. Since GacH and SccH are LtaS homologues, it is possible that the incorporated GroP is derived from phosphatidylglycerol yielding *sn*-Gro-1-P residues. To test this hypothesis, GroP was liberated from purified GAC by alkaline hydrolysis and separated from the polysaccharide by SEC. As explained in detail by Kennedy *et al* ^32^ for GroP-modified membrane oligosaccharides from *E. coli*, if GAC is modified by *sn*-Gro-1-P, alkaline hydrolysis of the phosphodiester bond should result in the formation of two cyclic intermediate compounds, Gro-1-cyclic phosphate and Gro-2-cyclic phosphate which further break up to a mixture of *sn*-Gro-1-P and Gro-2-P ^32^. If GAC is modified by *sn*-Gro-3-P, alkaline hydrolysis would yield a mixture of *sn*-Gro-3-P and Gro-2-P ^32^, whereas a phosphodiester of Gro-2-P would give a mixture of all three phosphates ^32^. Following alkaline hydrolysis the bulk of the carbohydrate still elutes in the void volume of the SEC column (Supplementary Fig. 10). However, the phosphate-containing fractions corresponding to the hydrolyzed GroP now elute in the inclusion volume (Supplementary Fig. 10). The enantiomeric composition of the GroP preparation was determined using a combination of LC-MS and an enzymatic method (see below). LC-MS revealed the presence of two GroP isomers, of approximately equal proportions, with LC retention times and major high molecular weight ions consistent with standard *sn*-Gro-1-P and Gro-2-P (Fig. 4f–4h, Supplementary Fig. 11). To resolve whether *sn*-Gro-3-P or *sn*-Gro-1-P is the substituent, the recovered GroP was characterized further by enzymatic analysis using a commercially available *sn*-Gro-3-P assay kit. Under reaction conditions in which 500 pmol of *sn*-Gro-3-P produced a robust enzymatic signal, incubation with either 500 pmol of *sn*-Gro-1-P or 500 pmol of the unknown Gro-P, recovered following alkaline hydrolysis, resulted in negligible activity (Supplementary Fig. 12). When a mixture containing 500 pmol of standard *sn*-Gro-3-P, and an equal amount of either *sn*-Gro-1-P or the unknown mixture of Gro-P isomers were tested, 85.8% and 90.0% of the activity detected with 500 pmol *sn*-Gro-3-P, alone, was found, demonstrating that the negative result using the unknown mixture, by itself, was not due to the presence of an unknown inhibitory compound in the GroP preparation. Taken together, our results indicate that GacH decorates GAC with *sn*-Gro-1-P, which is most probably derived from phosphatidylglycerol.

### NMR spectroscopy confirms the presence of GroP at the C6 hydroxyl group of GlcNAc side-chains

To unambiguously prove the incorporation of GroP in GAC, the polysaccharide isolated from WT GAS as described above was employed for NMR analysis (Fig. 5 a-g). ^1^H and ^13^C NMR spectra of GAC confirmed the presence of Rha and GlcNAc in a 1.7:1 ratio as determined by a chemical analysis of its sugar components. Although the material was heterogeneous, initial analysis revealed that it would be possible to analyze the purified GAC at different levels of detail. The major component identifiable at the highest level of intensity in a multiplicity-edited ^1^H,^13^C-HSQC NMR spectrum (Supplementary Fig. 13), acquired by non-uniform sampling at the 25% level of coverage facilitating enhanced resolution to resolve spectral overlap ^33^, revealed the ^1^H NMR chemical shifts in agreement with the following structure →3)-α-L-Rha*p*-(1→2)[β-D-Glc*p*NAc-(1→3)]-α-L-Rha*p*-(1→ as its repeating unit; ^7^ the ^1^*J*_HC_-based correlations facilitated identification of the resonances of the proton-carrying ^13^C nuclei.

**Fig. 5.**
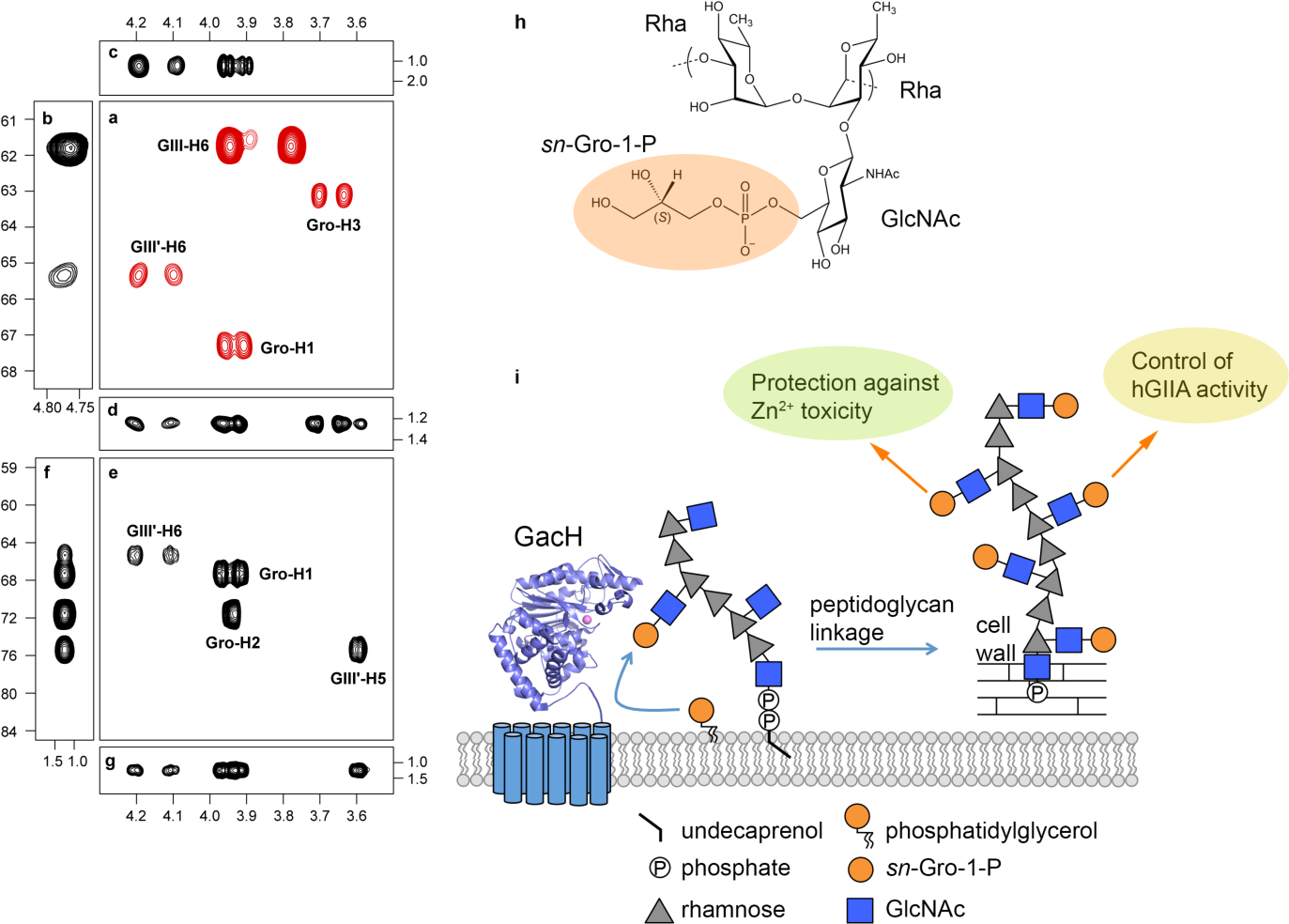
NMR analysis confirms presence of GroP on GlcNAc hydroxymethyl group of GAC. (**a**-**g**) Selected regions of NMR spectra of GAC. (**a**) Multiplicity-edited ^1^H,^13^C-HSQC in which methylene groups have opposite phase and are shown in red color, (**b**) ^1^H,^13^C-HSQC-TOCSY with an isotropic mixing time of 120 ms, (**c**) ^1^H,^13^C-HMBC with a mixing time of 90 ms, (**d**) ^1^H,^31^P-hetero-TOCSY with an isotropic mixing time of 80 ms, (**e**) ^1^H,^13^C-plane, (**f**) ^13^C, ^31^P-plane using a nominal ^n^*J*_CP_ value of 5 Hz, and (**g**) ^1^H, ^31^P-plane of a through-bond 3D ^1^H,^13^C,^13^P NMR experiment. Cross-peaks are annotated as GIII corresponding to the GlcNAc residue, GIIIʹ being the GroP-substituted GlcNAc residue and Gro as the glycerol residue. NMR chemical shifts of ^1^H (horizontal axis), ^13^C (left axis) and ^31^P (right axis and panel **f**) are given in ppm. (**h**) Schematic structure of the GAC repeating unit consisting of →3)-α-L-Rha*p*-(1→2)[β-D-Glc*p*NAc6*P*(*S*)Gro-(1→3)]-α-L-Rha*p*-(1→. (**i**) The mechanism and the roles of GroP cell wall modification in streptococci.

An array of 2D NMR experiments including ^1^H,^1^H-TOCSY, ^1^H,^1^H-NOESY, ^1^H,^13^C-HMBC and ^1^H,^13^C-HSQC-TOCSY led to the ^1^H and ^13^C NMR chemical shift assignments (Supplementary Table 2) and confirmed the structure of the trisaccharide repeating unit. A number of additional cross-peaks were visible at the second intensity level of the ^1^H,^13^C-HSQC spectrum, particularly in the spectral region of methylene groups, e.g. hydroxymethyl groups of hexopyranses ^34^. Besides the resonances from the hydroxymethyl group of the β-D-Glc*p*NAc residue at δ_H_ 3.78 and 3.94 correlated with a signal at δ_C_ 61.8, three more ^13^C signals together with corresponding ^1^H resonances were present at δ_C_ 63.1, 65.4 and 67.3 (Fig. 5a), readily identified from the multiplicity-editing applied in the experiment. In the ^13^C NMR spectrum two resonances at δ_C_ 67.3 and 71.6 were conspicuous in that they were split by 5.6 and 7.6 Hz, respectively. A ^13^C resonance observed as a doublet in this manner suggests scalar coupling to the NMR active nucleus ^31^P ^35^ and acquisition of a ^31^P NMR spectrum on GAC showed a prominent resonance at δ_P_ 1.2. The ^1^H,^1^H-TOCSY NMR spectrum at the second intensity level revealed chemical shift displacements in the region for anomeric resonances (Supplementary Fig. 14), and in the ^1^H,^13^C-HSQC-TOCSY spectrum two correlations from the anomeric proton resonance at δ_H_ ~4.76 were observed to δ_C_ 61.8 (C6 in β-D-Glc*p*NAc) and to δ_C_ 65.4 (Fig. 5b). Employing a ^1^H,^31^P-HMBC experiment revealed correlations to protons of the latter ^13^C at δ_H_4.10 and 4.19 as well as to δ_H_ 3.91 and 3.96, the ^13^C nucleus of which resonates at δ_C_ 67.3 (Fig. 5c). Moreover, a ^1^H,^31^P-hetero-TOCSY experiment showed in addition to the just described proton correlations further correlations to δ_H_ 3.59, 3.64 and 3.70 (Fig. 5d). Using the arsenal of 2D NMR experiments the resonances at the second level deviating from the parent structure were assigned (Supplementary Table 2). Thus, the GAC is partially substituted by a GroP residue at O6 of the side-chain β-D-Glc*p*NAc residue; based on integration of the cross-peaks for the anomeric resonances in the ^1^H,^13^C-HSQC NMR spectrum, the GAC preparation carries GroP groups to ~30 % of the GlcNAc residues. To validate the above results, a triple-resonance ^1^H,^13^C,^31^P NMR experiment based on through-bond ^1^*J*_HC_ as well as ^2^*J*_CP_ and ^3^*J*_CP_ correlations ^36^ was carried out. The 3D NMR experiment revealed the ^1^H NMR chemical shifts of H5ʹ and the two H6ʹ protons of the β-D-Glc*p*NAc residue, as well as the two H1ʹ protons and H2ʹ of the Gro residue all correlated to ^13^C nuclei (Fig. 5e). The ^13^C NMR chemical shifts of C5ʹ and C6ʹ of the β-D-Glc*p*NAc residue as well as C1ʹ and C2ʹ of the Gro residue all correlated to the ^31^P nucleus (Fig. 5f), and the above protons correlated to the ^31^P nucleus (Fig. 5g). Taking into considerations the GacH-mediated mechanism of GAC modification by GroP as well as the biochemical experiments carried out herein, the substituent at O6 of β-D-Glc*p*NAc is an *sn*-Gro-1-P group (Fig. 5h).

## Discussion

In Gram-positive bacteria, the common theme of most peptidoglycan-attached carbohydrate-based polymers is the presence of negatively charged groups in the repeating units ^3^. For example, canonical and non-canonical WTAs and Group B carbohydrate of *Streptococcus agalactiae* contain phosphodiester groups in the repeating units, peptidoglycan of *B. subtilis* grown in phosphate limiting conditions is decorated with a teichuronic acid containing glucuronic acid, and secondary cell wall carbohydrates of *Bacillus anthracis* are modified with pyruvyl groups ^2–4^. In bacteria lacking WTA such as the human pathogens GAS and *S. mutans*, it has been proposed that other polyanionic structures, such as LTA, fulfill similar functions during the bacterial cell cycle ^4,37^. Previous detailed studies using immunochemical methods, composition and linkage analyses and NMR methods deduced chemical structures of peptidoglycan-attached SRPs from both streptococcal species ^7,8,38-42^, but none identified anionic groups in these structures, except one overlooked study in which the presence of glycerol and phosphate in GAC has been detected ^43^. However, it was proposed that this GroP is part of the phosphodiester linkage connecting GAC to the N-acetylmuramic acid of peptidoglycan ^43^. Similarly, a number of reports identified substantial concentrations of phosphate in SRPs isolated from *Streptococcus sanguis*, *Streptococcus gallolyticus*, *Streptococcus dysgalactiae*, *Streptococcus sobrinus* serotype *d* and *S. mutans* serotype *f* ^30,42,44,45^. These phosphate-rich polymers were either disregarded as contamination with LTA ^30^, or further analyzed using ^1^H NMR or ^13^C NMR methods ^7,8,40-42,46,47^ that do not directly detect phosphoryl moieties in polysaccharides. With our report, we unambiguously confirm that SRPs of GAS and *S. mutans* are in fact polyanionic glycopolymers through decoration of their respective glycan side-chains with GroP (Fig. 5i).

We identified and structurally characterized a new family member of GroP transferase enzyme, GacH, which is required for GAC modification with GroP in GAS. GacH homologues are present in the SRP biosynthetic loci of many streptococci suggesting that a large proportion of streptococcal species produce SRPs where glycosyl side-chains are decorated with GroP similar to our observations for *S. mutans*. GacH is predicted to be an extracellular protein anchored in the cytoplasmic membrane. It belongs to the alkaline phosphatase superfamily of which two GroP transferases involved in LTA synthesis, LtaS and LtaP, have been biochemically and structurally characterized to date ^25-27,29^. LtaS and LtaP are membrane proteins that use the membrane lipid phosphatidylglycerol as the GroP donor for the transfer reaction ^48,49^. Our structural analysis of GacH in complex with GroP indicates that the enzyme uses the catalytic T530 residue to participate in the formation of a GroP-enzyme intermediate. This corresponds to the structure of LtaS, where the GroP molecule is complexed in the active site in which a threonine residue functions as a nucleophile in phosphatidylglycerol hydrolysis ^25–27^. The observation that the GroP in GAC is the *sn*-Gro-1-P enantiomer, strongly suggests that GacH uses phosphatidylglycerol as its donor substrate, similar to LtaS (Fig. 5i). Thus our data propose a catalytic mechanism in which GacH binds phosphatidylglycerol and T530 functions as a catalytic nucleophile to attack the GroP head group and cleave the phosphodiester bond. Diacylglycerol is released, leaving a covalent GroP-T530 intermediate. Next, the C6 hydroxyl group of the GAC GlcNAc target is deprotonated and then attacks the GroP-T530 enzyme, forming the GroP-GlcNAc product and returning the enzyme to the apo state.

In Gram-positive bacteria, the tailoring modification of WTA and LTA with D-alanine provides resistance against antibiotics, cationic antimicrobial peptides and small bactericidal enzymes including hGIIA, and promotes Mg^2+^ ion scavenging ^1–3^. It has been assumed that incorporation of positively charged D-alanine into teichoic acids decrease negative bacterial surface charge resulting in reduced initial binding of cationic antimicrobial peptides to the bacterial surface due to ionic repulsion ^50,51^. Our study demonstrates that addition of the negatively charged GroP group to SRPs protects streptococci from zinc toxicity but also renders bacteria more sensitive to hGIIA activity. A large body of published evidence indicates that phagocytic cells utilize Zn^2+^ intoxication to suppress the intracellular survival of bacteria ^52^. The mechanism of microbial susceptibility to zinc toxicity is mediated by extracellular competition of Zn^2+^ for Mn^2+^ transport and thereby mediating toxicity by impairing Mn^2+^ acquisition ^53^. Accordingly, the phenotypes of our mutants deficient in the GroP modifications and the GlcNAc side-chains could be explained either by “trapping” of Zn^2+^ in the WT cell wall as a consequence of zinc binding to negatively-charged GroP, or the increased Mn^2+^-binding capacity of GroP-modified bacterial cell wall which has been proposed to act as the conduit for the trafficking of mono- and divalent cations to the membrane ^2^.

Charge-dependent mechanisms are likely also underlying the increased hGIIA susceptibility of GAS and *S. mutans* expressing the GroP-modified SRPs. hGIIA is a highly cationic enzyme that catalyzes the hydrolysis of bacterial phosphatidylglycerol ^54–56^, ultimately leading to bacterial death. It has been suggested that traversal of this bactericidal enzyme across the Gram-positive cell wall to reach the plasma membrane is charge dependent because the absence of positively-charged D-alanine modifications in LTA and WTA severely compromises *S. aureus* survival when challenged with hGIIA ^56,57^. This phenotype was attributed to increased binding of hGIIA to the cell surface or modified permeability of the cell envelope. Similarly, the GacH/SccH-dependent GroP modifications on SRPs are required for hGIIA to exert its bactericidal effect against GAS and *S. mutans*, respectively. We have previously demonstrated that loss of the GlcNAc GAC side-chain strongly hampers trafficking of hGIIA through the GAS cell wall, with a minor contribution of reduced hGIIA binding to the cell surface ^21^. Since GroP-modifications were also lost in the GlcNAc side-chain deficient mutant, 5448Δ*gacI*, described in this study, we now assume that the mechanisms of the hGIIA-dependent phenotype are similar in the *gacI* and *gacH* mutants.

Another very important aspect of our study is the identification of a novel, potentially antigenic, epitope on the surface of streptococcal bacteria. GAS is a major human pathogen, and is associated with numerous diseases ranging from minor skin and throat infections such as impetigo and pharyngitis to life-threatening invasive diseases such as scarlet fever, streptococcal toxic syndrome and rapidly progressing deep-tissue infections, necrotizing fasciitis (i.e. the “flesh eating disease”), cellulitis and erysipelas ^58^. GAS infections are also responsible for post-infectious autoimmune syndrome, rheumatic fever (RF) and its sequelae, rheumatic heart disease (RHD) ^58^. Invasive GAS infections are difficult to treat with antibiotics and a GAS vaccine is urgently needed to combat this neglected disease. GAC is an attractive candidate for GAS vaccine due to its conserved expression in all GAS serotypes and the absence of the constitutive component of GAC, Rha, in humans ^16,17^. However, it has been proposed that the GAC GlcNAc side-chain can elicit the cross-reactive antibodies relevant to the pathogenesis of RF and RHD ^59–61^. Moreover, persistence of anti-GAC and anti-GlcNAc antibodies is a marker of poor prognosis in RHD ^60,62^. These clinical associations and the lack of understanding of the pathogenesis of GAS post-infectious RHD have hampered progress in the development of GAC-based vaccines against GAS. However, the GAC GlcNAc decorated with GroP might be a feasible candidate for GAS vaccine development because modified GlcNAc represents a unique epitope, that is absent from human tissues. Thus, our study has implications for design of a safe and effective vaccine against this important human pathogen for which a vaccine is not yet available. Finally, our work provides a framework for structure-function investigations of cell wall modifications in streptococci.

## Methods

### Bacterial strains, growth conditions and media

All plasmids, strains and primers used in this study are listed in Supplementary Tables 3 and 4. Streptococcal strains used in this study were the M1-serotype GAS strains, 5448 ^14^, 2221 ^63^ and 5005 ^63^, and *c* serotype *S. mutans* Xc ^64^. GAS and *S. mutans* strains were grown in Todd-Hewitt broth supplemented with 1% yeast extract (THY) without aeration at 37 °C. *S. mutans* plates were grown with 5% CO_2_. For hGIIA-mediated killing experiments *S. mutans* strains were grown in Todd-Hewitt broth without yeast extract and with 5% CO_2_. *E. coli* strains were grown in Lysogeny Broth (LB) medium or on LB agar plates at 37 °C. When required, antibiotics were included at the following concentrations: ampicillin at 100 μg/mL for *E. coli*; streptomycin at 100 μg/mL for *E. coli*; erythromycin (Erm) at 500 μg/mL for *E. coli* and 5 μg/mL for GAS and *S. mutans*; chloramphenicol (CAT) at 10 µg/mL for *E. coli* and 2 µg/mL for GAS and *S. mutans*; spectinomycin at 200 μg/mL for *E. coli*, 100 μg/mL for GAS and 500 μg/mL for *S. mutans*.

To identify GAS genes providing resistance against zinc toxicity, we used chemically defined medium based on the formulation of RPMI 1640 medium ^65^, which mirrors the amino acid composition of the van de Rijn and Kessler formulation ^66^. This RPMI 1640 (without glucose) (Gibco) is supplemented with nucleobases guanine, adenine and uracil at a concentration of 25 µg/mL each, as well as D-glucose at a final concentration of 0.5% w/v and HEPES at 50 mM. Necessary vitamins for GAS growth are provided by 100X BME Vitamins (Sigma B6891). The final solution (mRPMI) is pH 7.4 and capable of supporting GAS growth without additional supplements.

### Genetic manipulations

#### DNA techniques

Plasmids were transformed into GAS and *S. mutans* by electroporation or natural transformation as described previously ^9,67^. Chromosomal DNA was purified from GAS and *S. mutans* as described ^68^. All constructs were confirmed by sequencing analysis (Eurofins MWG Operon and Macrogen).

#### Genetic manipulation of GAS 5005 and 2221

For construction of the 5005Δ*gacH* and 2221Δ*gacH* strains, 5005 chromosomal DNA was used as a template for amplification of two DNA fragments using two primers pairs: 5005-f/gacHdel-r and gacHdel-f/5005-r. Primer gacHdel-f is complementary to primer gacHdel-r. The two gel-purified PCR products containing complementary ends were mixed and amplified using a PCR overlap method ^69^ with primer pair 5005-f/5005-r to create the deletion of *gacH*. The PCR product was digested with BamHI and XhoI and ligated into BamHI/SalI-digested plasmid pBBL740. The integrational plasmid pBBL740 does not have a replication origin that is functional in GAS, so the plasmid can be maintained only by integrating into the GAS chromosome through homologous recombination. The plasmid was designated pBBL740Δ*gacH*. The resulting plasmid was transformed into 5005 and 2221, and CAT resistant colonies were selected on THY agar plates. Five randomly selected colonies, that had the first crossover, were grown in liquid THY without CAT for ≥5 serial passages. Several potential double crossover mutants were selected as previously described ^70^. The deletion in each mutant was confirmed by PCR sequencing of the loci.

To construct the plasmid for *in cis* complementation of the 5005Δ*gacH* mutant, 5005 chromosomal DNA was used as a template for amplification of a wild-type copy of *gacH* using the primer pair 5005-f/5005-r. The PCR products were digested with BamHI and XhoI, and cloned in pBBL740 previously digested with the respective enzymes. The plasmid was designated pBBL740*gacH*. The plasmid was transformed into the 5005Δ*gacH* strain, and CAT resistant colonies were selected on THY agar plates. Double crossover mutants were selected as described above. Selected mutants were confirmed by PCR sequencing, yielding strain *5005*Δ*gacH:gacH*.

#### Genetic manipulation of GAS 5448

For construction of the 5448Δ*gacH* strains, GAS 5448 chromosomal DNA was used to amplify up and downstream regions flanking *gacH* using the following primer pairs: 5448-f/5448CAT-r and 5448CAT-f/5448-r. Primers 5448CAT-f and 5448CAT-r contain 25 bp extensions complementary to the CAT resistance cassette. Up- and downstream were fused to the CAT cassette using 5448-f/5448-r, digested with XhoI and HindIII and ligated into XhoI/HindIII-digested plasmid pHY304, yielding plasmid pHY304Δ*gacH*. After transformation in electrocompetent GAS 5448, transformed colonies were selected in THY containing Erm at 30 °C. After confirmation by PCR, transformed colonies were shifted to the non-permissive temperature of 37 °C to allow plasmid integration. Serial passage at 30 °C in the absence of antibiotic enabled occurrence of double cross-over events, yielding 5448Δ*gacH*, which were identified by screening for Erm sensitivity and CAT resistance. Deletion of *gacH* was confirmed by PCR.

To complement 5448Δ*gacH*, we created an expression plasmid p*gacH*_erm. *GacH* was amplified from GAS 5448 chromosomal DNA using primer pair gacH-EcoRI-f/gacH-BglII-r, digested using EcoRI/BglII, and ligated into EcoRI/BglII-digested pDCerm. p*gacH*_erm was transformed into the electrocompetent 5448Δ*gacH* and selected for Erm resistance on THY agar plates. Transformation was confirmed by PCR, yielding strain *5448*Δ*gacH:*p*gacH*.

*Genetic manipulation of S. mutans* Xc. For construction of the *sccH* deletion mutant (SMUΔ*sccH*), *S. mutans* Xc chromosomal DNA was used to amplify up and downstream regions flanking using the following primer pairs: sccH-f/sccH-erm-r and sccH-erm-f /sccH-r. Primers sccH-erm-f and sccH-erm-r contain 25 bp extensions complementary to the Erm resistance cassette. Up and downstream PCR fragments were mixed with the Erm cassette and amplified as a single PCR fragment using primer pair sccH-f/sccH-r. The *sccH* knockout construct was transformed into *S. mutans* as described previously ^9^. Erm resistant single colonies were picked and checked for deletion of *sccH* and integration of Erm cassette by PCR using primer pair: sccH-c-f/sccH-c-r, resulting in SMUΔ*sccH*. For complementation, *sccH* and *gacH* were amplified from *S. mutans Xc* and GAS 5448 chromosomal DNA, respectively, using primer pairs sccH-EcoRI-f/sccH-BglII-r and gacH-EcoRI-f/gacH-BglII-r. The PCR products were digested with EcoRI/BglII, and ligated into EcoRI/BglII-digested pDC123 vector, yielding p*sccH* and p*gacH*_cm, respectively. The plasmids were transformed into SMUΔ*sccH* as described ^9^. CAT resistant single colonies were picked and checked for presence of p*sccH* or p*gacH*_cm by PCR, yielding strains SMUΔ*sccH:*p*sccH* and SMUΔ*sccH:*p*gacH*, respectively.

#### Construction of the plasmids for *E. coli* expression of gacH

To create a vector for expression of extracellular domain of GacH, the gene was amplified from 5005 chromosomal DNA using the primer pair gacH-NcoI-f and gacH-XhoI-r. The PCR product was digested with NcoI and XhoI, and ligated into NcoI/XhoI-digested pET-NT vector. The resultant plasmid, pETGacH, contained *gacH* fused at the N-terminus with a His-tag followed by a TEV protease recognition site.

### Protein expression and purification

For expression and purification of eGacH, *E. coli* Rosetta (DE3) cells carrying the respective plasmid were grown to an OD_600_ of 0.4-0.6 and induced with 0.25 mM isopropyl β-D-1-thiogalactopyranoside (IPTG) at 18 °C for approximately 16 hrs. The cells were lysed in 20 mM Tris-HCl pH 7.5, 300 mM NaCl with two passes through a microfluidizer cell disrupter. The soluble fractions were purified by Ni-NTA chromatography with washes of 20 mM Tris-HCl pH 7.5, 300 mM NaCl and 20 mM Tris-HCl pH 7.5, 300 mM NaCl, 10 mM imidazole, and elution with 20 mM Tris-HCl pH 7.5, 300 mM NaCl, 250 mM imidazole. The eluate was dialyzed into 20 mM Tris-HCl pH 7.5, 300 mM NaCl in the presence of TEV protease (1 mg per 20 mg of protein). The dialyzed sample was reapplied to a Ni-NTA column equilibrated in 20 mM Tris-HCl pH 7.5, 300 mM NaCl to remove the cleaved His-tag and any uncleaved protein from the sample. eGacH was further purified by size exclusion chromatography (SEC) on a Superdex 200 column in 20 mM HEPES pH 7.5, 100 mM NaCl, with monitoring for protein elution at 280 nm. Fractions collected during elution from the column were analyzed for purity by SDS-PAGE and concentrated to approximately 10 mg/mL.

For expression of seleno-methionine labeled eGacH, *E. coli* Rosetta (DE3) carrying eGacH was grown in LB at 37 °C until an optical density at 600 nm of approximately 0.5 was obtained. The bacteria were centrifuged and resuspended in M9 minimal media supplemented with seleno-methionine. After a significant increase in optical density, protein expression was induced with 0.25 mM IPTG, and the cultures were grown at 16 °C for approximately 16 hrs. Seleno-methionine labeled eGacH was purified as described above.

### Crystallization, data collection and structure solution

eGacH crystallization conditions were initially screened using the JCSG Suites I–IV screens (Qiagen) at a protein concentration of 9 mg/mL by hanging drop vapor diffusion method. Crystals of Se-Met-substituted eGacH were grown in 0.1 M HEPES pH 7.5, 10% PEG8000, 8% ethylene glycol. Crystals were transferred into crystallization solution supplemented with 20% ethylene glycol and flash frozen in liquid nitrogen. The data were collected at APS 22-ID at a wavelength of 0.9793 Å. Crystals of GroP•eGacH complex were obtained using crystallization solution containing 0.2 M calcium acetate, 0.1 M MES pH 6.0, 20% PEG8000. *sn*-glycerol-1-phosphate (Sigma Aldrich) was mixed with eGacH at 10 mM prior to crystallization. Initial crystals of GroP•eGacH complex belonged to the same crystal form as apo GacH, however, crystals of different morphology grew epitaxially after several days. These crystals displayed better diffraction and were used for structure determination of GroP•eGacH complex. Crystals were cryoprotected in crystallization solution supplemented with 10 mM *sn*-glycerol-1-phosphate and 20% ethylene glycol and vitrified in liquid nitrogen. The data were collected at SSRL BL9-2 at a wavelength of 0.97946 Å.

All data were processed and scaled using *XDS* and *XSCALE* ^71^. The structure of eGacH was solved by Se single-wavelength anomalous diffraction method. Se atoms positions were determined using HySS module in *PHENIX* ^72,73^. The structure was solved using AutoSol wizard in *PHENIX* ^74^. The model was completed using *Coot* ^75^ and refined using *phenix.refine* ^76^. The final structure has two eGacH molecules in the asymmetric unit containing residues 444–822.

The structure of GroP•eGacH complex was solved by molecular replacement using *Phaser* ^77^ and the dimer of apo eGacH as a search model. The model was adjusted using *Coot* and refined using *phenix.refine*. Difference electron density corresponding to GroP molecules was readily identified after refinement. GroP molecules were modeled using *Coot*. The geometric restraints for GroP were generated using Grade Web Server (http://grade.globalphasing.org) (Global Phasing). The last several rounds of refinement were performed using 19 translation/libration/screw (TLS) groups, which were identified by *PHENIX* ^78^.

The structures were validated using *Coot*, MolProbity ^79^ and wwPDB Validation Service (https://validate.wwpdb.org) ^80^. Statistics for data collection, refinement, and model quality are listed in Supplementary Table 5. The structure factors and coordinates were deposited to the Protein Data Bank with accession codes 5U9Z (apo eGacH) and 6DGM (GroP•eGacH complex). Structure figures were generated using PyMOL v1.8.0.3 ^81^.

### Isolation of cell wall

Cell wall was isolated from exponential phase cultures by the SDS-boiling procedure as described for *S. pneumoniae* ^82^. Purified cell wall samples were lyophilized and used for carbohydrate composition analysis, phosphate and glycerol assays. Cell wall isolated from GAS 5005 was used for purification of GAC for *sn*-glycerol-1-phosphate identification and NMR analysis.

### GAC purification

GAC was released from the cell wall by sequential digestion with mutanolysin (Sigma Aldrich) and recombinant PlyC amidase ^19^, and partially purified by a combination of SEC and ion-exchange chromatography. Mutanolysin digests contained 5 mg/mL of cell wall suspension in 0.1 M sodium acetate, pH 5.5, 2 mM CaCl_2_ and 5 U/mL mutanolysin. Following overnight incubation at 37 °C, soluble polysaccharide was separated from the cell wall by centrifugation at 13,000 x g, 10 min. Acetone (-20 °C) was added to a final concentration of 80% and the polysaccharide was allowed to precipitate overnight at -20 °C. The precipitate was sedimented (5,000 x g, 20 min), dried briefly under nitrogen gas and redissolved in 0.1 M Tris-Cl, pH 7.4 PlyC (50 µg/mL) was added to the GAC sample and the reaction was incubated overnight at 37 °C. Following PlyC digestion, GAC was recovered by acetone precipitation, as described above, redissolved in a small volume of 0.2 N acetic acid and chromatographed on a 25 mL column of BioGel P10 equilibrated in 0.2 N acetic acid. Fractions (1.5 mL) were collected and monitored for carbohydrate by the anthrone assay. Fractions containing GAC (eluting near the void volume of the column) were combined, concentrated by spin column centrifugation (3,000 MW cutoff filter) and desalted by several rounds of dilution with water and centrifugation. After desalting, GAC was loaded onto an 18 mL column of DEAE-Sephacel. The column was eluted with a 100 mL gradient of NaCl (0-1 M). Fractions were analyzed for carbohydrate by the anthrone assay and phosphate by the malachite green assay following digestion with 70% perchloric acid (see below). Fractions containing peaks of carbohydrate were combined, concentrated by spin column (3,000 MW cut off) and lyophilized.

### Anthrone assay

Total carbohydrate content was determined by a minor modification of the anthrone procedure. Reactions containing 0.08 mL of aqueous sample and water were prepared in Safe-Lock 1.5 mL Eppendorf Tubes. Anthrone reagent (0.2% anthrone, by weight, dissolved in concentrated H_2_SO_4_) was rapidly added, mixed thoroughly, capped tightly and heated to 100 °C, 10 min. The samples were cooled in water (room temperature) and the absorbance at 580 nm was recorded. GAC concentration was estimated using an L-Rha standard curve.

### Phosphate assay

Approximately 1.5 mg of cell wall material isolated from GAS was dissolved in 400 µL H_2_O and 8 µg/mL PlyC, and incubated at 37 °C, rotating for approximately 16 hrs. Additional PlyC was added and incubated for a further 4-6 hrs. To liberate SCC from *S. mutans* cell wall, 1.5 mg of cell wall material isolated from *S. mutans* were incubated 24 h with 1.5 U/mL mutanolysin in 400 µL of 0.1 M sodium acetate, pH 5.5, 2 mM CaCl_2_. The samples were incubated at 100 °C for 20 min and centrifuged for 5 min at maximum speed in a table top centrifuge. The supernatant was transferred to a new micro-centrifuge tube and incubated with 2 N HCl at 100 °C for 2 hrs. The samples were neutralized with NaOH, in the presence of 62.5 mM HEPES pH 7.5. To 100 µL of acid hydrolyzed sample, 2 µL of 1 U/µL alkaline phosphatase (Thermo Fisher) and 10 µL 10 x alkaline phosphatase buffer was added and incubated at 37 °C, rotating, overnight. Released phosphate was measured using the Pi ColorLock Gold kit (Innova Biosciences), according to the manufacturer’s protocol.

During GAC purification on BioGel P10 and DEAE-Sephacel total phosphate content was determined by the malachite green method following digestion with perchloric acid. Fractions containing 10 to 80 µL were heated to 110 °C with 40 µL 70% perchloric acid (Fisher Scientific) in 13 x 100 borosilicate disposable culture tubes for 1 h. The reactions were diluted to 160 µL with water and 100 µL was transferred to a flat-bottom 96-well culture plate. Malachite Green reagent (0.2 mL) was added and the absorbance at 620 nm was read after 10 min at room temperature. Malachite Green reagent contained 1 vol 4.2% ammonium molybdate tetrahydrate (by weight) in 4 M HCl, 3 vol 0.045% malachite green (by weight) in water and 0.01% Tween 20.

### Glycerol assay

Samples for glycerol measurement were prepared as described for the phosphate assay but were not digested with alkaline phosphatase. Instead glycerol concentration was measured using the Glycerol Colorimetric assay kit (Cayman Chemical) according to the manufacturer’s protocol.

### Carbohydrate composition analysis

Carbohydrate composition analysis was performed at the Complex Carbohydrate Research Center (Athens, GA) by combined gas chromatography/mass spectrometry (GC/MS) of the per-O-trimethylsilyl derivatives of the monosaccharide methyl glycosides produced from the sample by acidic methanolysis as described previously ^83^.

### Identification of the stereochemistry of the GroP moiety of GAC

The stereochemistry of the GroP moiety attached to GAC was determined by a chemo-enzymatic method following release of GroP by alkaline hydrolysis as described by Kennedy *et al.* ^32^ using the Amplite^TM^ Fluorimetric *sn*-Glycerol-3-Phosphate (Gro-3-P) Assay Kit (AAT Bioquest). GAC was released from cell wall by sequential digestion with mutanolysin hydrolase and PlyC amidase, and partially purified by SEC on BioGel P10 and ion exchange chromatography on DEAE-Sephacel, as described above. GroP was liberated from the GAC by alkaline hydrolysis (0.5 M NaOH, 100 °C, 1 h), neutralized with acetic acid and recovered from the inclusion volume following SEC on BioGel P10. Column fractions containing GroP were identified by HPLC/mass spectrometry (LC-MS) and fractions containing GroP were combined, concentrated by rotary evaporation (30 °C, under reduced pressure) and desalted on BioGel P2. Column fractions containing GroP were combined, lyophilized and analyzed with the Gro-3-P Assay Kit based on the production of hydrogen peroxide in the *sn*-Gro-3-P oxidase-mediated enzyme coupled reaction. Reactions were conducted at room temperature for 10 to 20 min in solid black 96 well plates in a total volume of 0.1 ml. Excitation was at 540 nm and fluorescence at 590 nm was measured.

The fractions containing GroP were analyzed by LC-MS using a Q Exactive mass spectrometer and an Ultimate 3000 ultra high performance liquid chromatography system (Thermo Fisher Scientific). Chromatographic separation was achieved using a silica-based SeQuant ZIC-pHILIC column (2.1 mm × 150 mm, 5 µm, Merck, Germany) with elution buffers consisting of (A) 20 mM (NH_4_)_2_CO_3_ with 0.1% NH_4_OH in H_2_O and (B) acetonitrile. The column temperature was maintained at 40 °C, and the flow rate was set to 150 µL/min. Mass spectrometric detection was performed by electrospray ionization in negative ionization mode with source voltage maintained at 3.0 kV. The capillary temperature, sheath gas flow and auxiliary gas flow were set at 275 °C, 40 arb and 15 arb units, respectively. Full-scan MS spectra (mass range m/z 75 to 1000) were acquired with resolution R = 70,000 and AGC target 1e6.

### Identification of hGIIA-resistant GAS transposon mutants

The GAS M1T1 5448 *Krmit* transposon mutant library ^14^ was grown to mid-log phase (OD_600_ = 0.4). 1 x 10^5^ CFU were subjected to 27.5 µg/mL recombinant hGIIA ^84^ in triplicate and incubated for 1 h at 37 °C. Samples were plated on THY agar plates supplemented with kanamycin. The position of the transposon insertion of resistant colonies was determined as described previously ^85^.

### hGIIA susceptibility assay

hGIIA susceptibility experiments were performed as described previously ^21^. In short, mid-log suspensions (OD_600_ = 0.4), of GAS and *S. mutans* were diluted 1,000 times in HEPES solution (20 mM HEPES, 2 mM Ca^2+^, 1% BSA [pH 7.4]) and 10 µL was added to sterile round-bottom 96 well plates in triplicates. Recombinant hGIIA was serially diluted in HEPES solution and 10 µL aliquots were added to bacteria-containing wells. Samples were incubated for 2 hrs at 37 °C (for GAS without CO_2_, for *S. mutans* with 5% CO_2_), PBS was added and samples were 10-fold serially diluted for quantification on agar plates. Survival rate was calculated as Survival (% of inoculum) = (counted CFU * 100) / CFU count of original.

### Determination of selective metal concentrations

The Zn^2+^ sensitive gene deletion mutant 5448Δ*czcD* ^24^ was used to find the target concentration of Zn^2+^. Briefly, colonies of strains 5448 WT and 5448Δ*czcD* were scraped from THY agar plates and resuspended in PBS. After washing in PBS, the strains were adjusted to OD_600_ =1. These cultures were used to inoculate freshly prepared mRPMI containing varying concentrations of Zn^2+^ to starting OD_600_ = 0.05 in a 96-well plate. Growth at 37 °C was monitored at OD_595_ every 15 min using the BMG Fluostar plate reader.

### Tn-seq library screen for Zn^2+^ sensitivity

The 5448 *Krmit* Tn-seq library at T_0_ generation ^14^ was thawed, inoculated into 150 mL prewarmed THY broth containing 300 µg/mL kanamycin and grown at 37 °C for 6 hrs. After 6 hrs growth, the culture (T_1_) was centrifuged at 4,000 x *g* for 15 min at 4 °C and the pellet resuspended in 32.5 mL saline. Freshly prepared mRPMI or mRPMI containing 10 µM or 20 µM Zn^2+^ was inoculated with 500 µL culture into 39.5 mL media, creating a 1:20 fold inoculation. These T_2_ cultures were then grown at 37 °C for exactly 6 hrs, at which point 2 mL of these cultures were inoculated again into 38 mL of freshly prepared mRPMI alone or mRPMI containing 10 µM or 20 µM Zn^2+^. The remaining 38 mL of T_2_ culture was harvested by centrifugation at 4,000 x *g* for 10 min at 4 °C and pellets stored at -20 °C for later DNA extraction. Cultures were grown for a further 6 hrs, at which point T_3_ cultures were harvested by centrifugation at 4,000 x *g* for 10 min at 4 °C and pellets stored at -20 °C.

Tn-seq *Krmit* transposon insertion tags were prepared from the cell pellets as previously described ^15,86^. Briefly, genomic DNA was prepared using the MasterPure complete DNA and RNA purification kit (Epicentre) and treated with *MmeI* and the calf intestinal phosphatase (NEB) before ligation to 12 distinct Tn-seq adapters for sample multiplexing during massively parallel sequencing ^15,86^. Insertion tags were produced through a 22-cycle PCR ^15^ using the ligation mixtures and primers oKrmitTNseq2 and AdapterPCR ^15,86^. After quality control with the Bioanalyzer instrument (Agilent), the libraries of *Krmit* insertion tags were sequenced (50-nt single end reads) on an Illumina HiSeq 1500 in the Institute for Bioscience and Biotechnology Research (IBBR) Sequencing Core at the University of Maryland, College Park. Tn-seq read datasets were analyzed (quality, filtering, trimming, alignment, visualization) as previously described ^15,86^ using the M1T1 5448 genome as reference for read alignments. The ratios of mutant abundance comparing the output to input mutant pools were calculated as a fold change for each GAS gene using the DEseq2 and EdgeR pipelines ^86–88^. Illumina sequencing reads from the Tn-seq analysis were deposited in the NCBI Sequence Read Archive (SRA) under the accession number SRP150081.

### Drop test assays

Strains 5448 WT, 5448Δ*gacI*, 5448Δ*gacI:gacI*, 5448Δ*gacH*, 5448Δ*gacH*:p*gacH*, *S. mutans* WT, SMUΔ*sccH*, SMUΔ*sccH:*p*sccH* and SMUΔ*sccH:*p*gacH* were grown in THY to mid-exponential growth phase, adjusted to OD_600_ = 0.6 and serial diluted. 5 µL were spotted onto THY agar plates containing varying concentrations of Zn^2+^ (ZnSO_4_⃅7H_2_O). Plates were incubated at 37 °C overnight and photographed. Drop tests are representative of biological replicates performed on at least 3 separate occasions.

### NMR spectroscopy

The NMR spectra were recorded on a Bruker AVANCE III 700 MHz equipped with a 5 mm TCI Z-Gradient Cryoprobe (^1^H/^13^C/^15^N) and dual receivers and a Bruker AVANCE II 600 MHz spectrometer equipped with a 5 mm TXI inverse Z-Gradient ^1^H/D-^31^P/^13^C. The ^1^H and ^13^C NMR chemical shift assignments of the polysaccharide material were carried out in D_2_O solution (99.96 %) at 323.2 K unless otherwise stated. Chemical shifts are reported in ppm using internal sodium 3-trimethylsilyl-(2,2,3,3-^2^H_4_)-propanoate (TSP, *δ*_H_ 0.00 ppm), external 1,4-dioxane in D_2_O (*δ*_C_ 67.40 ppm) and 2 % H_3_PO_4_ in D_2_O (*δ*_P_ 0.00 ppm) as reference. The ^1^H,^1^H-TOCSY experiments (dipsi2ph) were recorded with mixing times of 10, 30, 60, 90 and 120 ms. The ^1^H,^1^H-NOESY experiments ^89^ were collected with mixing times of 100 and 200 ms. A uniform and non-uniform sampling (50 and 25 % NUS) were used for the multiplicity-edited ^1^H,^13^C-HSQC experiments ^90^ employing an echo/antiecho-TPPI gradient selection with and without decoupling during the acquisition. The 2D ^1^H,^13^C-HSQC-TOCSY were acquired using MLEV17 for homonuclear Hartman-Hahn mixing, an echo/antiecho-TPPI gradient selection with decoupling during acquisition and mixing times of 20, 40, 80 and 120 ms. The 2D ^1^H,^31^P-Hetero-TOCSY experiments ^91^ were collected using a DIPSI2 sequence with mixing times of 10, 20, 30, 50 and 80 ms. The 2D ^1^H,^31^P-HMBC experiments were recorded using an echo/antiecho gradient selection and mixing times of 25, 50 and 90 ms. The 3D ^1^H,^13^C,^31^P ^36^ spectra were obtained using echo/antiecho gradient selection and constant time in *t*_2_ with a nominal value of^n^*J*_CP_ of 5 Hz and without multiplicity selection. The spectra were processed and analyzed using TopSpin 4.0.1 software (Bruker BioSpin).

### Bioinformatics analysis

The TOPCONS (http://topcons.net/) ^92^ web server was employed to predict trans-membrane regions of GacH. Homology detection and structure prediction were performed by the HHpred server (https://toolkit.tuebingen.mpg.de/#/tools/hhpred) ^93^. To construct the GacH phylogenetic tree the homologues of full-length GacH (M5005_Spy_0609) were retrieved using blastp with an E-value cutoff of 1e^-70^. In addition, sequences were filtered based on a minimal identity of 33%, similarity of 73%, and having 11 predicted transmembrane helixes. Of all species expressing *gacH* homologues, *ltaS* homologues were retrieved using blastp and GAS *ltaS* (M5005_Spy_0622) as reference. As representatives of the LtaS and LtaP clades, five sequences of Listeria were selected that express both enzymes ^27^. All protein sequences were aligned using MUSCLE ^94^. The phylogenetic tree was build using MEGA6 ^95^ and the Maximum Likelihood method based on the JTT matrix-based model ^95^. For the phylogenetic tree with the extracellular domains of GacH and LtaS homologues, extracellular domains were predicted using http://www.cbs.dtu.dk/services/TMHMM/.

## Statistical analysis

Unless otherwise indicated, statistical analysis was carried out on pooled data from at least three independent biological repeats. Quantitative data was analyzed using the paired Student’s t-test. A 2-way ANOVA with Bonferroni multiple comparison test was used to compare multiple groups. A *P*-value equal to or less that 0.05 was considered statistically significant.

## Acknowledgements

This work was supported by the Center of Biomedical Research Excellence (COBRE) Pilot Grant (to NK, KVK and JSR) supported by NIH grant P30GM110787 from the National Institute of General Medical Sciences (NIGMS), VIDI grant 91713303 from the Netherlands Organization for Scientific Research (NWO) (to NMvS and VPvH), the Swedish Research Council (no. 2013-4859 and 2017-03703) and The Knut and Alice Wallenberg Foundation (to GW), NIH grant P30GM110787 from the NIGMS and NIH grant 1S10OD021753 (to AJM), the National Health and Medical Research Council of Australia (to MJW), grants from CNRS, ANR (MNaims ANR-17-CE17-0012-01) and FRM (SPF20150934219) (to GL), NIH grant AI047928 from the National Institute of Allergy and Infectious Diseases (NIAID) (to KSM and YLB) and NIH grant AI094773 (to NES and ATB).

Carbohydrate composition analysis at the Complex Carbohydrate Research Center was supported by the Chemical Sciences, Geosciences and Biosciences Division, Office of Basic Energy Sciences, U.S. Department of Energy grant (DE-FG02-93ER20097) to Parastoo Azadi. Use of the Advanced Photon Source was supported by the U. S. Department of Energy, Office of Science, Office of Basic Energy Sciences, under Contract No. W-31-109-Eng-38 and NIH grants S10_RR25528 and S10_RR028976. Use of the Stanford Synchrotron Radiation Lightsource, SLAC National Accelerator Laboratory, is supported by the U.S. Department of Energy, Office of Science, Office of Basic Energy Sciences under Contract No. DE-AC02-76SF00515. The SSRL Structural Molecular Biology Program is supported by the DOE Office of Biological and Environmental Research, and by the NIH, NIGMS including P41GM103393. The contents of this publication are solely the responsibility of the authors and do not necessarily represent the official views of NIGMS or NIH.

## Author contributions

AR, PD, YLB, KSM, AGM, AJM, GL, MJW, JSR, KVK, GW, NMvS and NK designed the experiments. RJE, VPvH, AR, AT, JSR, KVK, GW and NK performed functional and biochemical experiments. KVK carried out X-ray crystallography and structure analysis. AR and GW performed NMR studies. PD and AJM performed MS analysis. VPvH and NK constructed plasmids and isolated mutants. RJE, VPvH, AR, PD, YLB, NMES, ATB, KSM, AGM, AJM, MJW, JSR, KVK, GW, NMvS and NK analyzed the data. NMvS and NK wrote the manuscript with contributions from all authors. All authors reviewed the results and approved the final version of the manuscript.

## References

1. Brown, S., Santa Maria, J.P., Jr. & Walker, S. Wall teichoic acids of Gram-positive bacteria. Annu. Rev. Microbiol. 67, 313–336 (2013).

2. Neuhaus, F.C. & Baddiley, J. A continuum of anionic charge: structures and functions of D-alanyl-teichoic acids in Gram-positive bacteria. Microbiol. Mol. Biol. Rev. 67, 686–723 (2003).

3. Weidenmaier, C. & Peschel, A. Teichoic acids and related cell-wall glycopolymers in Gram-positive physiology and host interactions. Nat. Rev. Microbiol. 6, 276–287 (2008).

4. Mistou, M.Y., Sutcliffe, I.C. & van Sorge, N.M. Bacterial glycobiology: rhamnose containing cell wall polysaccharides in Gram-positive bacteria. FEMS Microbiol. Rev. 40, 464–479 (2016).

5. McCarty, M. The lysis of group A hemolytic streptococci by extracellular enzymes of *Streptomyces albus*. II. Nature of the cellular substrate attacked by the lytic enzymes. J. Exp. Med. 96, 569–580 (1952).

6. Lancefield, R.C. A serological differentiation of human and other groups of hemolytic Streptococci. J. Exp. Med. 57, 571–595 (1933).

7. Huang, D.H., Rama Krishna, N. & Pritchard, D.G. Characterization of the group A streptococcal polysaccharide by two-dimensional^1^H-nuclear-magnetic-resonance spectroscopy. Carbohydr. Res. 155, 193–199 (1986).

8. St Michael, F. et al. Investigating the candidacy of the serotype specific rhamnan polysaccharide based glycoconjugates to prevent disease caused by the dental pathogen *Streptococcus mutans*. Glycoconj. J. 35, 53–64 (2018).

9. van der Beek, S.L. et al. GacA is essential for Group A Streptococcus and defines a new class of monomeric dTDP-4-dehydrorhamnose reductases (RmlD). Mol. Microbiol. 98, 946–962 (2015).

10. Tsuda, H., Yamashita, Y., Shibata, Y., Nakano, Y. & Koga, T. Genes involved in bacitracin resistance in *Streptococcus mutans*. Antimicrob. Agents Chemother. 46, 3756–3764 (2002).

11. De, A. et al. Deficiency of RgpG causes major defects in cell division and biofilm formation, and deficiency of LytR-CpsA-Psr family proteins leads to accumulation of cell wall antigens in culture medium by *Streptococcus mutans*. Appl. Environ. Microbiol. 83, e00928 (2017).

12. Nagata, E. et al. Serotype-specific polysaccharide of *Streptococcus mutans* contributes to infectivity in endocarditis. Oral. Microbiol. Immunol. 21, 420–423 (2006).

13. van Sorge, N.M. et al. The classical Lancefield antigen of Group A Streptococcus is a virulence determinant with implications for vaccine design. Cell Host Microbe 15, 729–740 (2014).

14. Le Breton, Y. et al. Essential genes in the core genome of the human pathogen *Streptococcus pyogenes*. Sci. Rep. 5, 9838 (2015).

15. Henningham, A. et al. Virulence role of the GlcNAc side chain of the Lancefield cell wall carbohydrate antigen in non-M1-serotype Group A Streptococcus. mBio 9, e02294 (2018).

16. Kabanova, A. et al. Evaluation of a Group A Streptococcus synthetic oligosaccharide as vaccine candidate. Vaccine 29, 104–114 (2010).

17. Sabharwal, H. et al. Group A streptococcus (GAS) carbohydrate as an immunogen for protection against GAS infection. J. Infect. Dis. 193, 129–135 (2006).

18. Shibata, Y., Yamashita, Y., Ozaki, K., Nakano, Y. & Koga, T. Expression and characterization of streptococcal *rgp* genes required for rhamnan synthesis in *Escherichia coli*. Infect. Immun. 70, 2891–2898 (2002).

19. Rush, J.S. et al. The molecular mechanism of N-acetylglucosamine side-chain attachment to the Lancefield group A carbohydrate in *Streptococcus pyogenes*. J. Biol. Chem. 292, 19441–19457 (2017).

20. Ozaki, K. et al. A novel mechanism for glucose side-chain formation in rhamnose glucose polysaccharide synthesis. FEBS Lett. 532, 159–163 (2002).

21. van Hensbergen, V.P. et al. Streptococcal Lancefield polysaccharides are critical cell wall determinants for human Group IIA secreted phospholipase A2 to exert its bactericidal effects. PLoS Pathog. 14, e1007348 (2018).

22. Weiss, J.P. Molecular determinants of bacterial sensitivity and resistance to mammalian Group IIA phospholipase A2. Biochim. Biophys. Acta. 1848, 3072–3077 (2015).

23. Graham, M.R. et al. Virulence control in group A Streptococcus by a two-component gene regulatory system: global expression profiling and *in vivo* infection modeling. Proc. Natl. Acad. Sci. USA 99, 13855–13860 (2002).

24. Ong, C.L., Gillen, C.M., Barnett, T.C., Walker, M.J. & McEwan, A.G. An antimicrobial role for zinc in innate immune defense against group A streptococcus. J. Infect. Dis. 209, 1500–1508 (2014).

25. Lu, D. et al. Structure-based mechanism of lipoteichoic acid synthesis by *Staphylococcus aureus* LtaS. Proc. Natl. Acad. Sci. USA 106, 1584–1589 (2009).

26. Schirner, K., Marles-Wright, J., Lewis, R.J. & Errington, J. Distinct and essential morphogenic functions for wall-and lipo-teichoic acids in *Bacillus subtilis*. EMBO J. 28, 830–842 (2009).

27. Campeotto, I. et al. Structural and mechanistic insight into the *Listeria monocytogenes* two-enzyme lipoteichoic acid synthesis system. J. Biol. Chem. 289, 28054–28069 (2014).

28. Schneewind, O. & Missiakas, D. Lipoteichoic acids, phosphate-containing polymers in the envelope of Gram-positive bacteria. J. Bacteriol. 196, 1133–1142 (2014).

29. Webb, A.J., Karatsa-Dodgson, M. & Grundling, A. Two-enzyme systems for glycolipid and polyglycerolphosphate lipoteichoic acid synthesis in *Listeria monocytogenes*. Mol. Microbiol. 74, 299–314 (2009).

30. Emdur, L.I., Saralkar, C., McHugh, J.G. & Chiu, T.H. Glycerolphosphate-containing cell wall polysaccharides from *Streptococcus sanguis*. J. Bacteriol. 120, 724–732 (1974).

31. Fischer, W., Laine, R.A. & Nakano, M. On the relationship between glycerophosphoglycolipids and lipoteichoic acids in Gram-positive bacteria. II. Structures of glycerophosphoglycolipids. Biochim. Biophys. Acta 528, 298–308 (1978).

32. Kennedy, E.P., Rumley, M.K., Schulman, H. & Van Golde, L.M. Identification of *sn* glycero-1-phosphate and phosphoethanolamine residues linked to the membrane derived oligosaccharides of *Escherichia coli*. J. Biol. Chem. 251, 4208–4213 (1976).

33. Orekhov, V.Y. & Jaravine, V.A. Analysis of non-uniformly sampled spectra with multi-dimensional decomposition. Prog. Nucl. Magn. Reson. Spectrosc. 59, 271–292 (2011).

34. Widmalm, G. A perspective on the primary and three-dimensional structures of carbohydrates. Carbohydr. Res. 378, 123–132 (2013).

35. Bernlind, C., Oscarson, S. & Widmalm, G. Synthesis, NMR, and conformational studies of methyl α-D-mannopyranoside 2-, 3-, 4-, and 6-monophosphates. Carbohydr. Res. 263, 173–180 (1994).

36. Marino, J.P. et al. Three-dimensional triple-resonance ^1^H, ^13^C, ^31^P experiment: sequential through-bond correlation of ribose protons and intervening phosphorus along the RNA oligonucleotide backbone. J. Am. Chem. Soc. 116, 6472–6473 (1994).

37. Caliot, E. et al. Role of the Group B antigen of *Streptococcus agalactiae*: a peptidoglycan-anchored polysaccharide involved in cell wall biogenesis. PLoS Pathog. 8, e1002756 (2012).

38. Coligan, J.E., Kindt, T.J. & Krause, R.M. Structure of the streptococcal groups A, A-variant and C carbohydrates. Immunochemistry 15, 755–760 (1978).

39. Mc, C.M. & Lancefield, R.C. Variation in the group-specific carbohydrate of group A streptococci. I. Immunochemical studies on the carbohydrates of variant strains. J. Exp. Med. 102, 11–28 (1955).

40. Pritchard, D.G., Coligan, J.E., Geckle, J.M. & Evanochko, W.T. High-resolution ^1^H-and ^13^C-n.m.r. spectra of the group A-variant streptococcal polysaccharide. Carbohydr. Res. 110, 315–319 (1982).

41. Pritchard, D.G., Gregory, R.L., Michalek, S.M. & McGhee, J.R. Characterization of the serotype e polysaccharide antigen of *Streptococcus mutans*. Mol. Immunol. 23, 141–145 (1986).

42. Pritchard, D.G., Michalek, S.M., McGhee, J.R. & Furner, R.L. Structure of the serotype f polysaccharide antigen of *Streptococcus mutans*. Carbohydr. Res. 166, 123–131 (1987).

43. Heymann, H., Manniello, J.M. & Barkulis, S.S. Structure of streptococcal cell walls. V. Phosphate esters in the walls of group A *Streptococcus pyogenes*. Biochem. Biophys. Res. Commun. 26, 486–491 (1967).

44. Pazur, J.H., Cepure, A., Kane, J.A. & Karakawa, W.W. Glycans from streptococcal cell walls: glycosyl-phosphoryl moieties as immunodominant groups in heteroglycans from group D and group L streptococci. Biochem. Biophys. Res. Commun. 43, 1421–1428 (1971).

45. Prakobphol, A., Linzer, R. & Genco, R.J. Purification and characterization of a rhamnose-containing cell wall antigen of *Streptococcus mutans* B13 (serotype d). Infect. Immun. 27, 150–157 (1980).

46. Czabanska, A., Holst, O. & Duda, K.A. Chemical structures of the secondary cell wall polymers (SCWPs) isolated from bovine mastitis *Streptococcus uberis*. Carbohydr. Res. 377, 58–62 (2013).

47. Neiwert, O., Holst, O. & Duda, K.A. Structural investigation of rhamnose-rich polysaccharides from *Streptococcus dysgalactiae* bovine mastitis isolate. Carbohydr. Res. 389, 192–195 (2014).

48. Karatsa-Dodgson, M., Wormann, M.E. & Grundling, A. *In vitro* analysis of the *Staphylococcus aureus* lipoteichoic acid synthase enzyme using fluorescently labeled lipids. J. Bacteriol. 192, 5341–5349 (2010).

49. Lequette, Y., Lanfroy, E., Cogez, V., Bohin, J.P. & Lacroix, J.M. Biosynthesis of osmoregulated periplasmic glucans in *Escherichia coli*: the membrane-bound and the soluble periplasmic phosphoglycerol transferases are encoded by the same gene. Microbiology 154, 476–483 (2008).

50. Peschel, A. et al. Inactivation of the *dlt* operon in *Staphylococcus aureus* confers sensitivity to defensins, protegrins, and other antimicrobial peptides. J. Biol. Chem. 274, 8405–8410 (1999).

51. Falagas, M.E., Rafailidis, P.I. & Matthaiou, D.K. Resistance to polymyxins: Mechanisms, frequency and treatment options. Drug Resist. Updat. 13, 132–138 (2010).

52. Djoko, K.Y., Ong, C.L., Walker, M.J. & McEwan, A.G. The role of copper and zinc toxicity in innate immune defense against bacterial pathogens. J. Biol. Chem. 290, 18954–18961 (2015).

53. McDevitt, C.A. et al. A molecular mechanism for bacterial susceptibility to zinc. PLoS Pathog. 7, e1002357 (2011).

54. Buckland, A.G. & Wilton, D.C. Inhibition of secreted phospholipases A2 by annexin V. Competition for anionic phospholipid interfaces allows an assessment of the relative interfacial affinities of secreted phospholipases A2. Biochim. Biophys. Acta 1391, 367–376 (1998).

55. Hsu, Y.H., Dumlao, D.S., Cao, J. & Dennis, E.A. Assessing phospholipase A2 activity toward cardiolipin by mass spectrometry. PLoS One 8, e59267 (2013).

56. Koprivnjak, T., Peschel, A., Gelb, M.H., Liang, N.S. & Weiss, J.P. Role of charge properties of bacterial envelope in bactericidal action of human group IIA phospholipase A2 against *Staphylococcus aureus*. J. Biol. Chem. 277, 47636–47644 (2002).

57. Hunt, C.L., Nauseef, W.M. & Weiss, J.P. Effect of D-alanylation of (lipo)teichoic acids of *Staphylococcus aureus* on host secretory phospholipase A2 action before and after phagocytosis by human neutrophils. J. Immunol. 176, 4987–4994 (2006).

58. Carapetis, J.R., Steer, A.C., Mulholland, E.K. & Weber, M. The global burden of group A streptococcal diseases. Lancet Infect. Dis. 5, 685–694 (2005).

59. Goldstein, I., Rebeyrotte, P., Parlebas, J. & Halpern, B. Isolation from heart valves of glycopeptides which share immunological properties with *Streptococcus haemolyticus* group A polysaccharides. Nature 219, 866–868 (1968).

60. Ayoub, E.M. & Dudding, B.A. Streptococcal group A carbohydrate antibody in rheumatic and nonrheumatic bacterial endocarditis. J. Lab. Clin. Med. 76, 322–332 (1970).

61. Kirvan, C.A., Swedo, S.E., Heuser, J.S. & Cunningham, M.W. Mimicry and autoantibody-mediated neuronal cell signaling in Sydenham chorea. Nat. Med. 9, 914–920 (2003).

62. Shulman, S.T. et al. Group A streptococcal pharyngitis serotype surveillance in North America, 2000-2002. Clin. Infect. Dis. 39, 325–332 (2004).

63. Sumby, P. et al. Evolutionary origin and emergence of a highly successful clone of serotype M1 group a Streptococcus involved multiple horizontal gene transfer events. J. Infect. Dis. 192, 771–782 (2005).

64. Koga, T., Asakawa, H., Okahashi, N. & Takahashi, I. Effect of subculturing on expression of a cell-surface protein antigen by *Streptococcus mutans*. J. Gen. Microbiol. 135, 3199–3207 (1989).

65. Moore, G.E., Gerner, R.E. & Franklin, H.A. Culture of normal human leukocytes. JAMA 199, 519–524 (1967).

66. van de Rijn, I. & Kessler, R.E. Growth characteristics of group A streptococci in a new chemically defined medium. Infect. Immun. 27, 444–448 (1980).

67. Hoff, J.S., DeWald, M., Moseley, S.L., Collins, C.M. & Voyich, J.M. SpyA, a C3-like ADP-ribosyltransferase, contributes to virulence in a mouse subcutaneous model of *Streptococcus pyogenes* infection. Infect. Immun. 79, 2404–2411 (2011).

68. Caparon, M.G. & Scott, J.R. Genetic manipulation of pathogenic streptococci. Methods Enzymol. 204, 556–586 (1991).

69. Horton, R.M., Hunt, H.D., Ho, S.N., Pullen, J.K. & Pease, L.R. Engineering hybrid genes without the use of restriction enzymes: gene splicing by overlap extension. Gene 77, 61–68 (1989).

70. Trevino, J., Liu, Z., Cao, T.N., Ramirez-Pena, E. & Sumby, P. RivR is a negative regulator of virulence factor expression in group A Streptococcus. Infect. Immun. 81, 364–372 (2013).

71. Kabsch, W. Xds. Acta Crystallogr. D Biol. Crystallogr. 66, 125–132 (2010).

72. Grosse-Kunstleve, R.W., & Adams, P.D. Substructure search procedures for macromolecular structures. Acta Crystallogr. D Biol. Crystallogr. 59, 1966–1973 (2003).

73. McCoy, A.J., Storoni, L.C. & Read, R.J. Simple algorithm for a maximum-likelihood SAD function. Acta Crystallogr. D Biol. Crystallogr. 60, 1220–1228 (2004).

74. Terwilliger, T.C. et al. Decision-making in structure solution using Bayesian estimates of map quality: the PHENIX AutoSol wizard. Acta Crystallogr. D Biol. Crystallogr. 65, 582–601 (2009).

75. Emsley, P., Lohkamp, B., Scott, W.G. & Cowtan, K. Features and development of *Coot*. Acta Crystallogr. D Biol. Crystallogr. 66, 486–501 (2010).

76. Afonine, P.V. et al. Towards automated crystallographic structure refinement with phenix.refine. Acta Crystallogr. D Biol. Crystallogr. 68, 352–367 (2012).

77. McCoy, A.J. et al. Phaser crystallographic software. J. Appl. Crystallogr. 40, 658–674 (2007).

78. Adams, P.D. et al. *PHENIX*: a comprehensive Python-based system for macromolecular structure solution. Acta Crystallogr. D Biol. Crystallogr. 66, 213–221 (2010).

79. Chen, V.B. et al. *MolProbity*: all-atom structure validation for macromolecular crystallography. Acta Crystallogr. D Biol. Crystallogr. 66, 12–21 (2010).

80. Gore, S. et al. Validation of structures in the Protein Data Bank. Structure 25, 1916–1927 (2017).

81. Schrodinger, LLC. The PyMOL Molecular Graphics System, Version 1.8.0.3. (2010).

82. Bui, N.K. et al. Isolation and analysis of cell wall components from *Streptococcus pneumoniae*. Anal. Biochem. 421, 657–666 (2012).

83. Santander, J. et al. Mechanisms of intrinsic resistance to antimicrobial peptides of *Edwardsiella ictaluri* and its influence on fish gut inflammation and virulence. Microbiology 159, 1471–1486 (2013).

84. Ghomashchi, F. et al. Preparation of the full set of recombinant mouse- and human-secreted phospholipases A2. Methods Enzymol. 583, 35–69 (2017).

85. Le Breton, Y. & McIver, K.S. Genetic manipulation of *Streptococcus pyogenes* (the Group A Streptococcus, GAS). Curr. Protoc. Microbiol. 30, Unit 9D 3 (2013).

86. Le Breton, Y. et al. Genome-wide discovery of novel M1T1 group A streptococcal determinants important for fitness and virulence during soft-tissue infection. PLoS Pathog. 13, e1006584 (2017).

87. Anders, S. & Huber, W. Differential expression analysis for sequence count data. Genome Biol. 11, R106 (2010).

88. Robinson, M.D., McCarthy, D.J. & Smyth, G.K. edgeR: a Bioconductor package fordifferential expression analysis of digital gene expression data. Bioinformatics 26, 139–140 (2010).

89. Wagner, R. & Berger, S. Gradient-selected NOESY-A fourfold reduction of the measurement time for the NOESY Experiment. J. Magn. Reson. A 123, 119–121 (1996).

90. Willker, W., Leibfritz, D., Kerssebaum, R. & Bermel, W. Gradient selection in inverse heteronuclear correlation spectroscopy. Magn. Reson. Chem. 31, 287–292 (1993).

91. Kellogg, G.W. Proton-detected hetero-TOCSY experiments with application to nucleic acids. J. Magn. Reson. 98, 176–182 (1992).

92. Tsirigos, K.D., Peters, C., Shu, N., Kall, L. & Elofsson, A. The TOPCONS web server for consensus prediction of membrane protein topology and signal peptides. Nucleic Acids Res. 43, W401–407 (2015).

93. Soding, J., Biegert, A. & Lupas, A.N. The HHpred interactive server for protein homology detection and structure prediction. Nucleic Acids Res. 33, W244–248 (2005).

94. Edgar, R.C. MUSCLE: multiple sequence alignment with high accuracy and high throughput. Nucleic Acids Res. 32, 1792–1797 (2004).

95. Tamura, K., Stecher, G., Peterson, D., Filipski, A. & Kumar, S. MEGA6: Molecular Evolutionary Genetics Analysis version 6.0. Mol. Biol. Evol. 30, 2725–2729 (2013).

