## Supplementary Information for "Discovery of glycerol phosphate modification on streptococcal rhamnose polysaccharides"

**Contents:**

Supplementary Figures 1-14
Supplementary Tables 1-5
Supplementary References

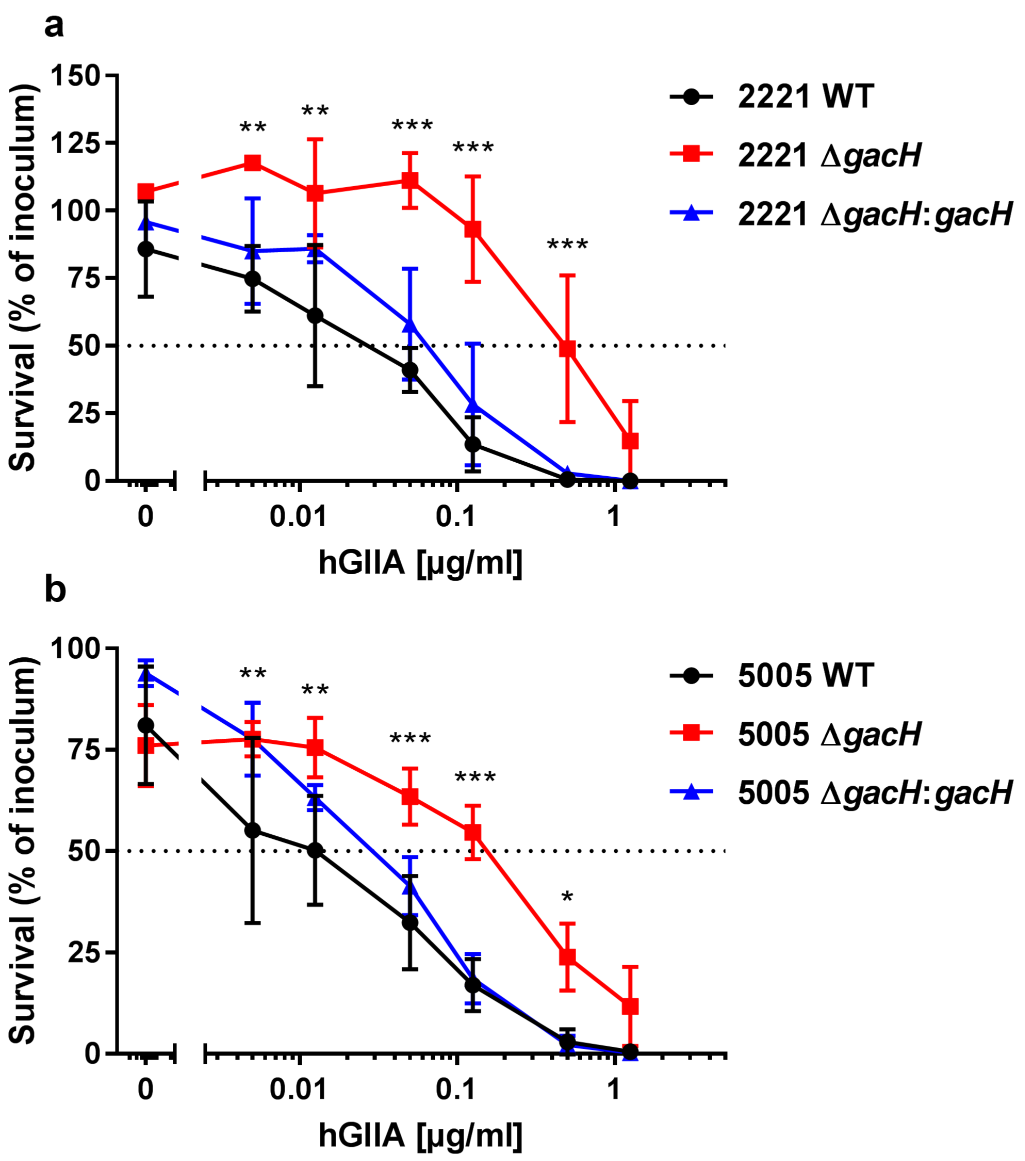


**Supplementary Fig. 1. Contribution of *gacH* to GAS hGIIA sensitivity in two different GAS strain backgrounds.**

**(a** and **b)** Deletion of *gacH* in (**a**) GAS 2221 and (**b**) GAS 5005 increases resistance to hGIIA-mediated killing in a concentration dependent manner as determined by CFU killing assay. Data represent mean ± standard deviation of three independent experiments. *, $p < 0.05$; **, $p < 0.01$; ***, $p < 0.001$.

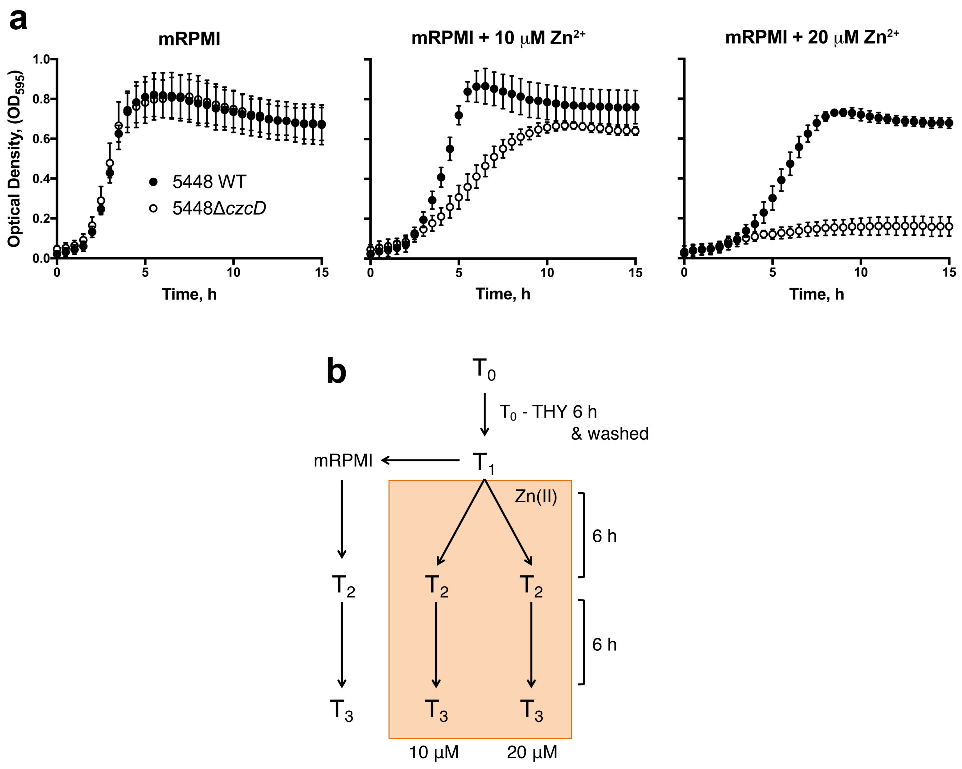


**Supplementary Fig. 2. Tn-seq library screen for $Zn^{2+}$ sensitivity.**

**(a)** Determination of $Zn^{2+}$ concentration for GAS Tn-seq experiment. Strains 5448 WT and 5448Δ*czcD* were grown in mRPMI alone, or containing 10 µM or 20 µM $Zn^{2+}$. Cultures were inoculated to starting $OD_{600}$ = 0.05 in 96-well plates and growth was monitored at $OD_{595}$ every 30 min. Graph represents mean ± standard deviation of three independent biological replicates. (**b)** Schematic of $Zn^{2+}$ toxicity Tn-seq experimentation. The GAS 5448 *Krmit* Tn-seq library at $T_0$ generation was inoculated into THY broth and grown at 37 °C for 6 hrs. After 6 hrs growth, the culture ($T_1$) was centrifuged and the pellet resuspended in saline. mRPMI or mRPMI containing 10 µM or 20 µM $Zn^{2+}$ was inoculated with bacterial culture creating a 1:20 fold inoculation. These $T_2$ cultures were then grown at 37 °C for 6 hrs, at which point 2 mL of these cultures were inoculated again into 38 mL of mRPMI or mRPMI containing 10 µM or 20 µM $Zn^{2+}$. The remaining 38 mL of $T_2$ culture was harvested by centrifugation and pellets used for DNA extraction. Cultures were grown for a further 6 hrs, at which point $T_3$ cultures were harvested by centrifugation and pellets used for DNA extraction.

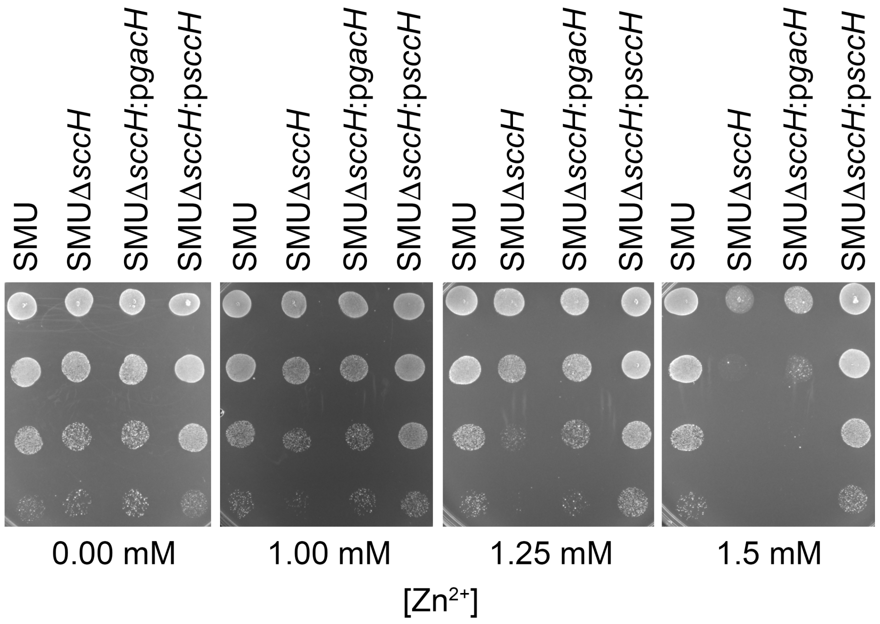


**Supplementary Fig. 3. $Zn^{2+}$ sensitivity analysis of *S. mutans* strains in drop test assay.**
*S. mutans* WT, Δ*sccH* and Δ*sccH* complemented with *gacH* or *sccH* were grown in THY to mid-exponential phase, adjusted to $OD_{600}$ = 0.6, serial diluted and 5 μL spotted onto THY agar plates containing varying concentrations of $Zn^{2+}$. The experiment was performed at least three times.

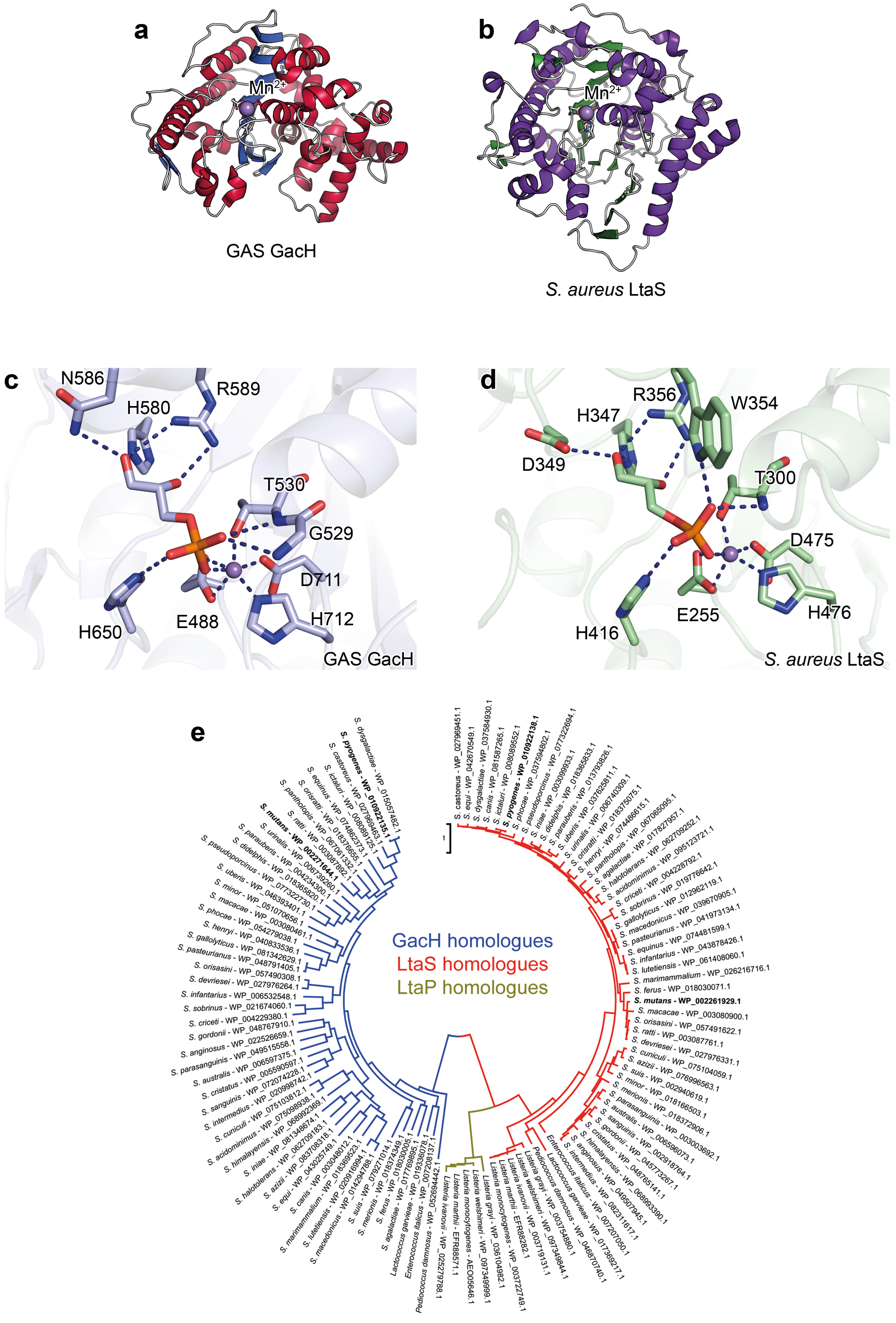


**Supplementary Fig. 4. Comparison of GacH and LtaS structures and phylogenetic analysis of the full-length GacH, LtaS, and LtaP enzymes.**

(**a**) Structure of apo eGacH viewing at the active site with the $Mn^{2+}$ ion shown as a violet sphere. (**b**) Structure of extracellular domain of *S. aureus* LtaS (PDB ID 2W5Q, [1]) shown in the same orientation as GacH. (**c**) A close-up view of the active site GacH crystal structure in complex with *sn*-Gro-1-P. (**d**) Comparison of the active site of LtaS (PDB ID 2W5T, [1]) with GroP in the active site. (**e**) Phylogenetic analysis by Maximum Likelihood method of the full-length GacH, LtaS, and LtaP enzymes. A phylogenetic tree was generated of 50 GacH (blue) and 50 LtaS (red) homologues from similar species. Five LtaP (yellow) homologues from *Listeria* species were selected to represent the LtaP clade. The GAS and *S. mutans* proteins analyzed in this study are indicated in bold. The phylogenetic tree is drawn to scale as indicated by the scale bar, with branch lengths measured in the number of substitutions per site.

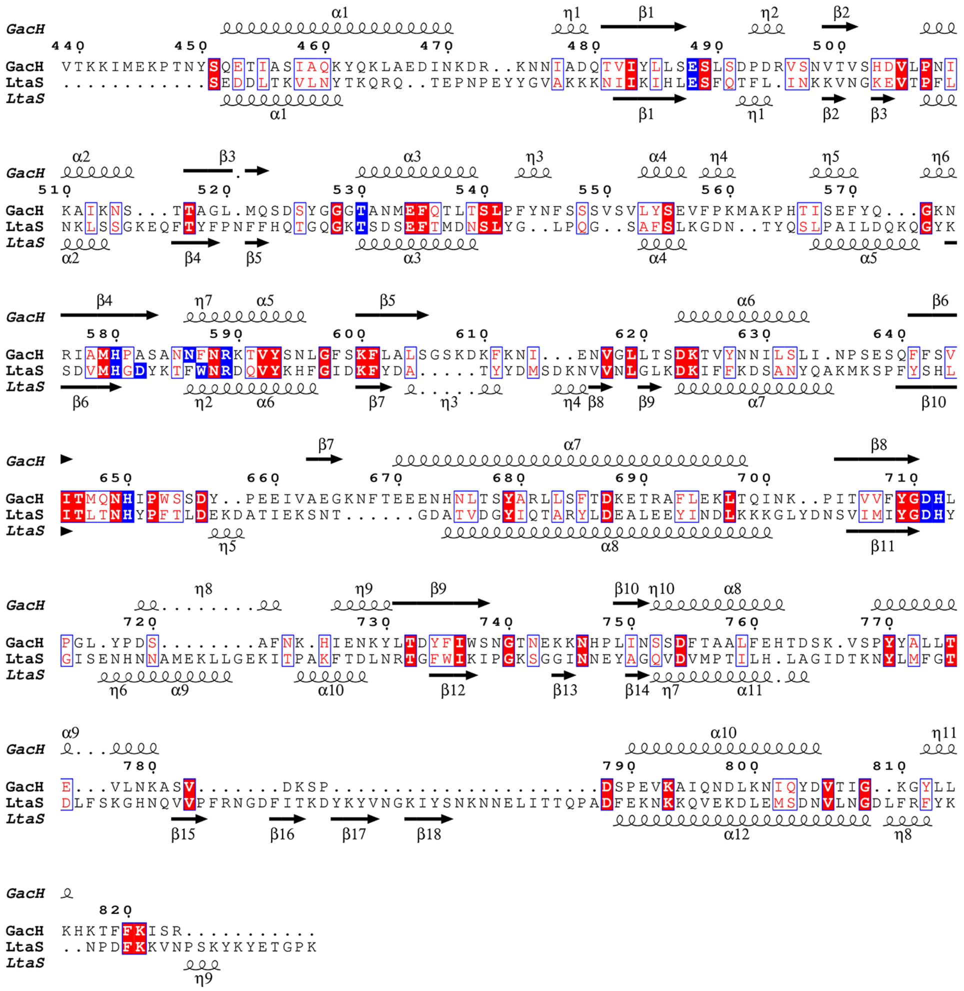

**Supplementary Fig. 5. Structure-based sequence alignment of extracellular domains of GAS GacH and *S. aureus* LtaS.**

Residues implicated in the catalytic cycle are highlighted in blue. The secondary structure elements are indicated above (PDB ID 5U9Z) and below (PDB ID 2W5Q, [1]) the alignment.

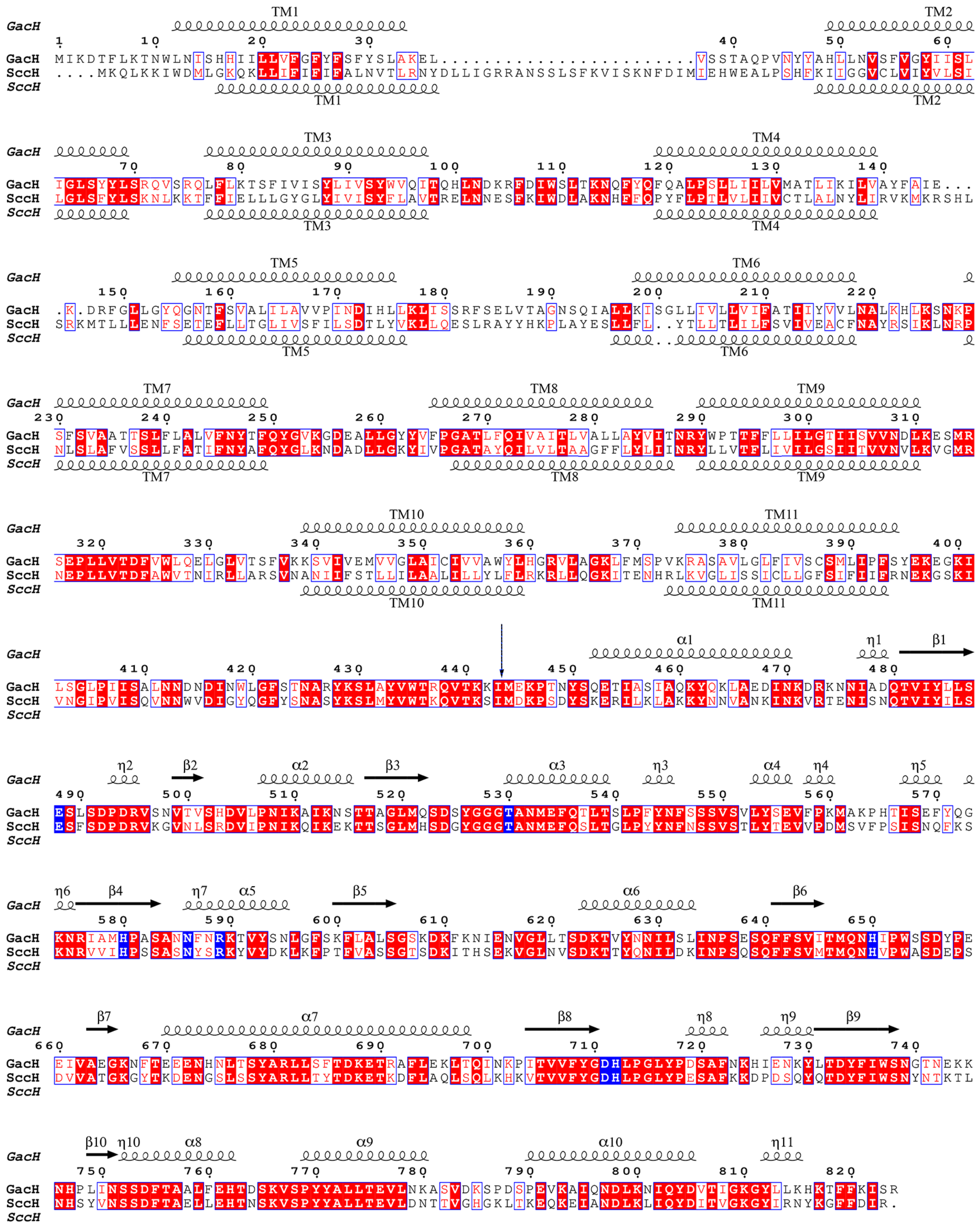

**Supplementary Fig. 6. Sequence alignment of GAS 5005 GacH and *S. mutans* Xc SccH.** Identical residues are highlighted in red. Residues involved in catalysis and GroP coordination are highlighted in blue. The predicted transmembrane helices for both proteins and the secondary structure elements of eGacH (PDB ID 5U9Z) are shown. Vertical arrow indicates the start of the construct used for crystallization of eGacH.

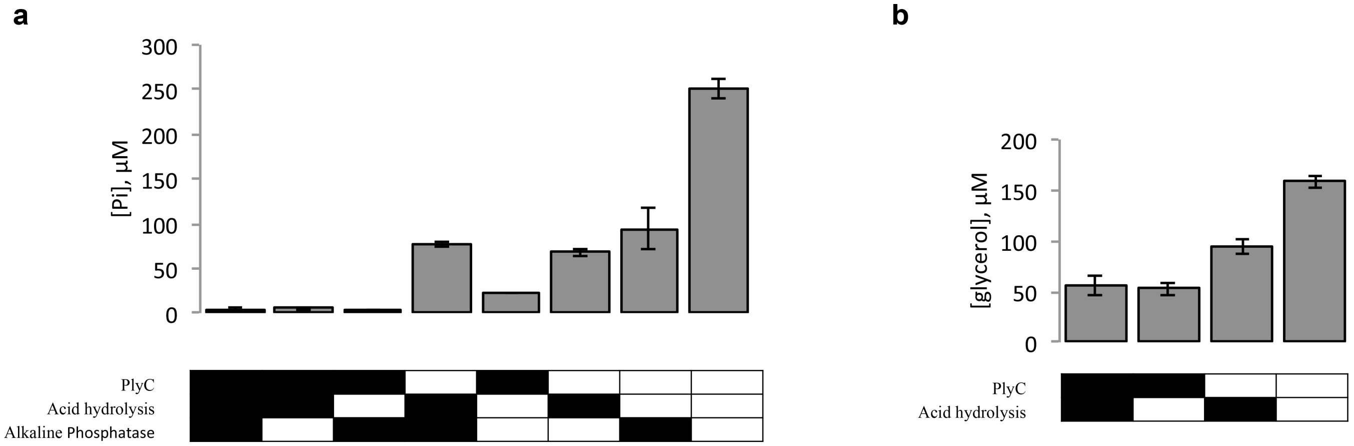


**Supplementary Fig. 7. Release of phosphate (Pi) and glycerol from cell wall isolated from GAS. (a** and **b)** Cell wall material isolated from GAS 5005 WT was subjected to a combination of PlyC digestion, acid hydrolysis and alkaline phosphatase digestion. The treatment conditions are indicated by black rectangles below the bar graph. (**a**) Pi concentration was measured for each condition using the malachite green assay. (**b)** Glycerol concentration was measured for each condition using the glycerol colorimetric assay kit. Data are the mean ± standard deviation of three biological replicates.

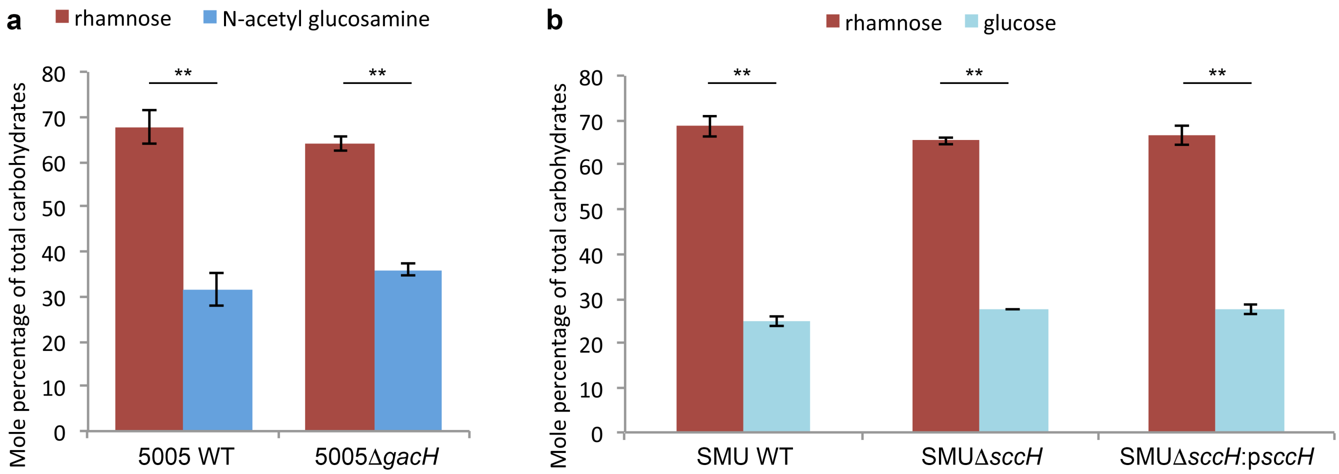


**Supplementary Fig. 8. The glycosyl composition of cell walls purified from the GAS and *S. mutans* strains.**

(**a** and **b)** Carbohydrate composition analysis of cell wall material. Rhamnose, N-acetylglucosamine and glucose mole percentage of total carbohydrate was determined by GC-MS for cell wall material isolated from (**a**) the GAS 5005 strains and (**b**) the *S. mutans* strains following methanolysis as described in Methods. Data are mean ± standard deviation of three replicates. **, $p < 0.01$.

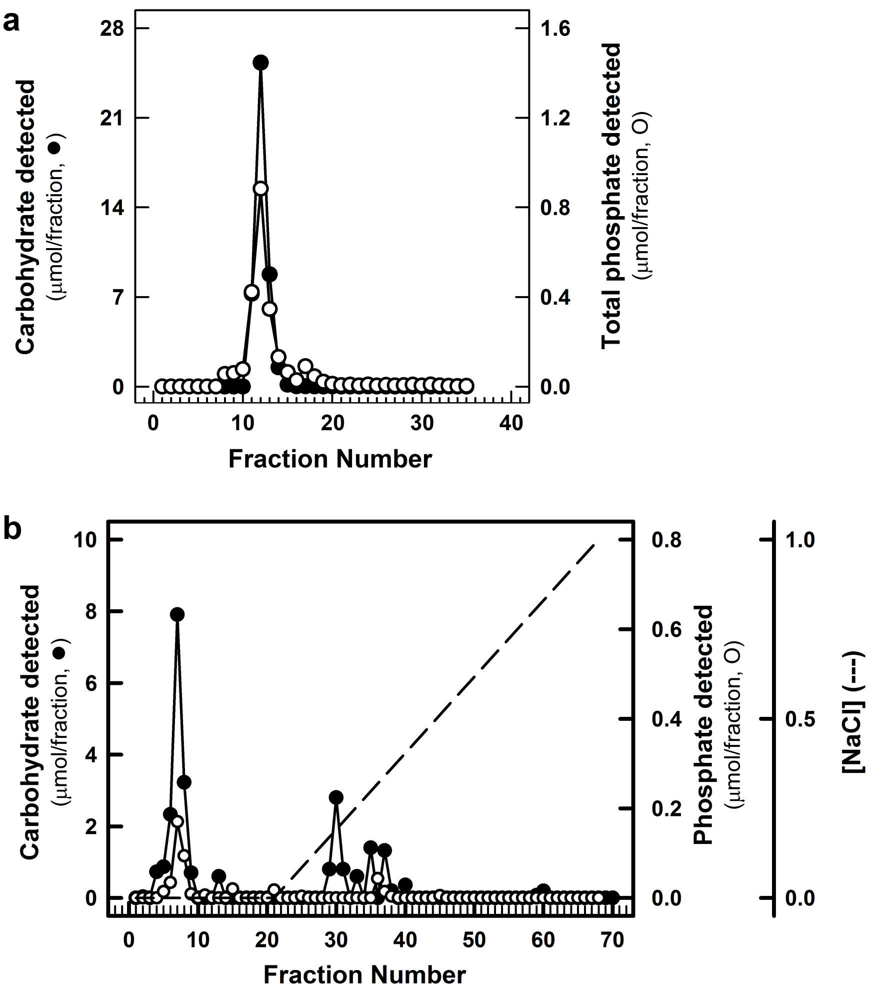


**Supplementary Fig. 9. GAC purification.**

**(a)** BioGel P10 chromatography of GAC isolated from GAS 5005 WT. Following PlyC and mutanolysin digestion, GAC was recovered by acetone precipitation, redissolved in 0.2 N acetic acid and chromatographed on a 25 mL column of BioGel P10 equilibrated in 0.2 N acetic acid. (**b)** DEAE-Sephacel elution profile of GAC purified from 5005Δ*gacH*. GAC was loaded onto an 18 mL column of DEAE-Sephacel. The column was eluted with a 100 mL gradient of NaCl (0-1 M). Fractions were analyzed for carbohydrate content by anthrone assay (●) and total phosphate content by malachite green assay following digestion with perchloric acid (O). Each experiment was performed at least three times.

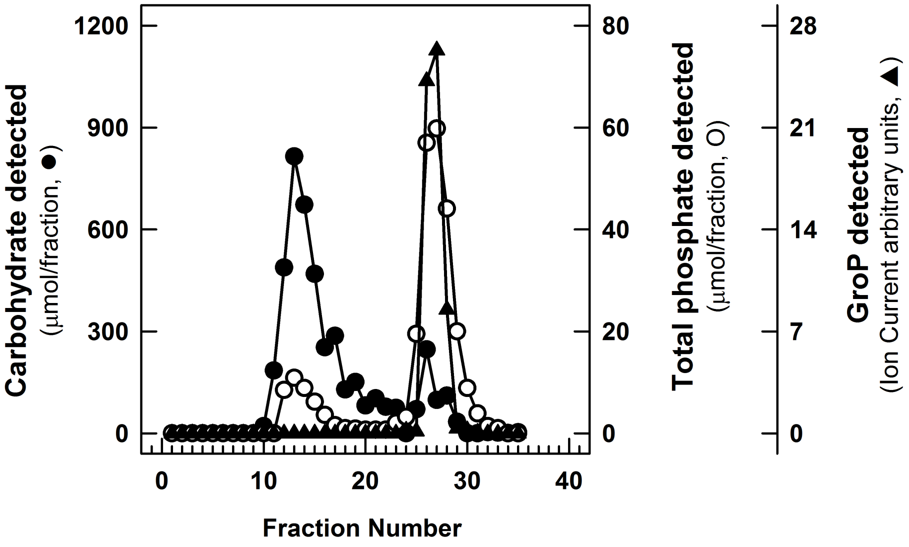


**Supplementary Fig. 10. BioGel P10 chromatography of DEAE-Sephacel bound GAC following alkaline hydrolysis.**

Fractions containing GAC from Fig. 4 e (T27-T36) were pooled, concentrated and desalted by spin column, hydrolyzed with alkali as described in Methods and chromatographed on a 25 mL column of BioGel P10, equilibrated in 0.2 N acetic acid. Fractions were analyzed for carbohydrate content by anthrone assay (●), total phosphate content by malachite green assay following digestion with perchloric acid (O) and for GroP by LC-MS (▲). The experiment was performed at least three times.

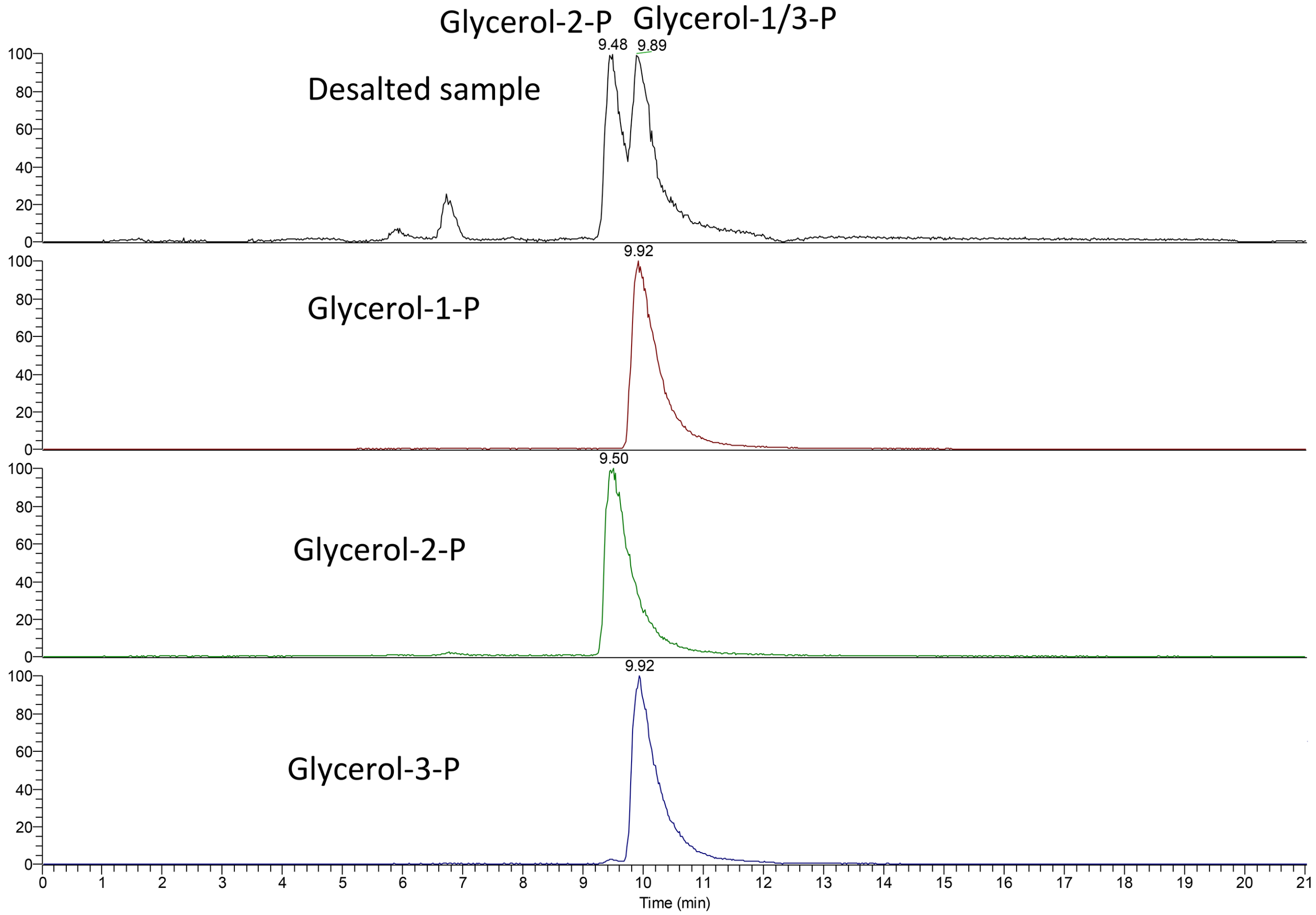


**Supplementary Fig. 11. Separation of GroP isomer standards and GroP recovered from GAC on a SeQuant ZIC-pHILIC column.**

The elution positions of GroP (top panel) recovered from GAC following alkaline hydrolysis and BioGel P10 chromatography (fractions T25-T30 containing phosphate from Supplementary Fig. 10), and standard *sn*-Gro-1-P, Gro-2-P and *sn*-Gro-3-P (bottom 3 panels) are shown. The GroP isomers were resolved using an Ultimate 3000 ultra high performance liquid chromatography system using a silica-based SeQuant ZIC-pHILIC column as described in Methods. The experiment was performed at least three times.

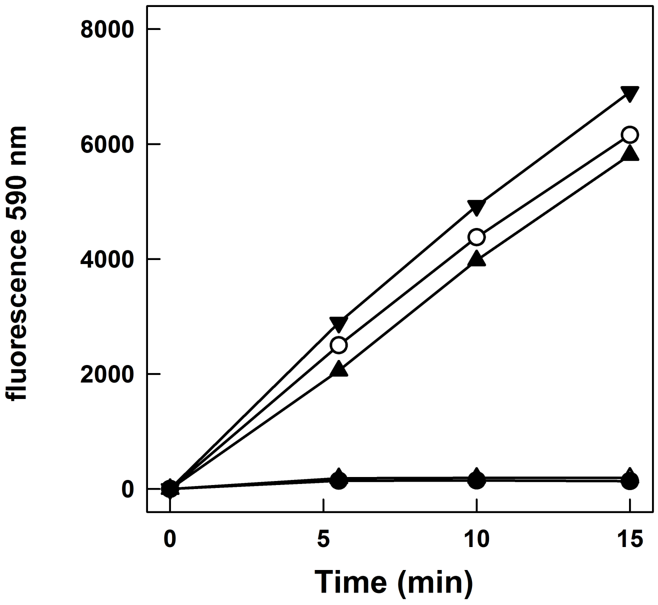


**Supplementary Fig. 12. Time course of *sn*-Gro-3-P detection by *sn*-Gro-3-P assay kit.**

*sn*-Gro-3-P was assayed using the Amplite$^{TM}$ Fluorimetric *sn*-Gro-3-P Assay Kit (AAT Bioquest) in the presence of either 500 pmol *sn*-Gro-3-P (▼), 500 pmol *sn*-Gro-1-P (△), 500 pmol unknown GroP (fractions T25-T30 containing GroP from Supplementary Fig. 10) (●), a mixture of 500 pmol *sn*-Gro-3-P and 500 pmol of unknown GroP (O) or a mixture of 500 pmol *sn*-Gro-3-P and 500 pmol *sn*-Gro-1-P (▲). Samples were analyzed with an excitation wavelength of 540 nm and emission detection at 590 nm. The experiment was performed at least three times.

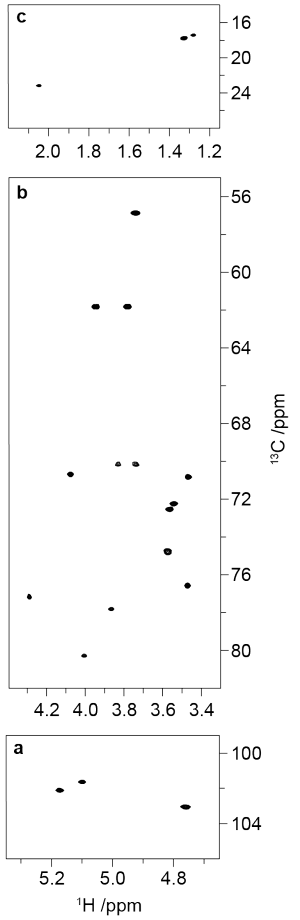


**Supplementary Fig. 13. Multiplicity-edited $^{1}$H,$^{13}$C-HSQC NMR spectrum of GAC recorded with non-uniform sampling of 25 %.**

(**a**-**c**) The resolution in the indirect dimension was enhanced by recording four times the number of points in the $^{13}$C fid to resolve the overlapped resonances **RI-4** and **RII-4**. The three sections show (**a**) the anomeric, (**b**) the ring and hydroxymethyl group and (**c**) the methyl group regions of the GAC repeating unit, respectively.

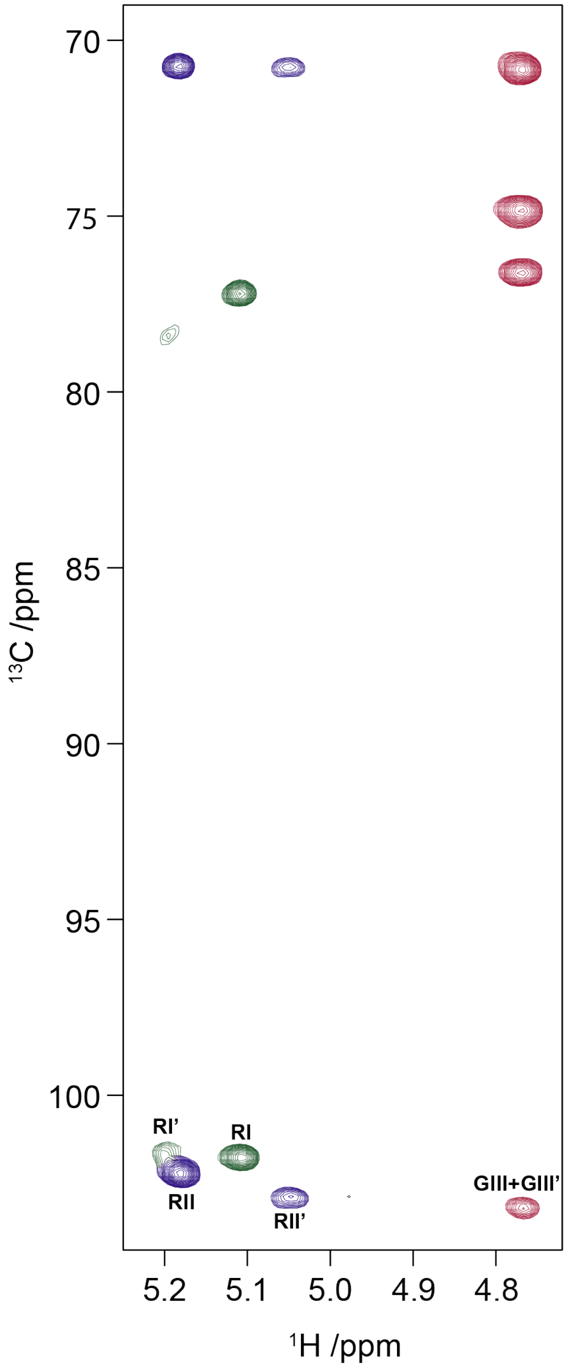


**Supplementary Fig. 14. Anomeric region of the $^{1}$H,$^{13}$C-HSQC-TOCSY spectrum of GAC recorded employing a 90 ms isotropic mixing time.**

Chemical shifts changes are shown for each anomeric resonance from the non-phosphorylated structure to the Gro*P*-carrying one, indicated by a prime. The $^{13}$C resonances of the two rhamnose ring systems (**RI** and **RII'**) in the phosphorylated form do not differ considerably to the non-phosphorylated counterparts unlike the anomeric ones.

**Supplementary Table 1.** Structural homologues of GacH.

| PDB ID | Z score | r.m.s.d. | Number of aligned residues | Total number of residues | Sequence identity, % | Protein function | Reference |
|---|---|---|---|---|---|---|---|
| 2w8d | 27.3 | 3.7 | 333 | 421 | 14 | lipoteichoic acid synthase | [2] |
| 2w5q | 26.9 | 3.9 | 333 | 424 | 16 | lipoteichoic acid synthase | [1] |
| 2w5r | 26.8 | 3.9 | 333 | 424 | 15 | lipoteichoic acid synthase | [1] |
| 2w5s | 26.6 | 3.8 | 332 | 424 | 16 | lipoteichoic acid synthase | [1] |
| 2w5t | 26.6 | 3.9 | 332 | 424 | 16 | lipoteichoic acid synthase | [1] |
| 4uop | 26.5 | 3.6 | 313 | 409 | 19 | lipoteichoic acid primase | [3] |
| 4uor | 26.4 | 3.9 | 332 | 418 | 15 | lipoteichoic acid synthase | [3] |
| 4uoo | 26.3 | 3.9 | 326 | 415 | 16 | lipoteichoic acid synthase | [3] |
| 3lxq | 25.7 | 4.3 | 310 | 409 | 15 | uncharacterized protein VP1736 | NP[a] |
| 2qzu | 18.6 | 3.9 | 279 | 465 | 11 | putative sulfatase yidJ | NP |
| 5i5d | 18.5 | 3.9 | 271 | 342 | 12 | inner membrane protein YejM | [4] |
| 6bnd | 18.4 | 3.4 | 257 | 335 | 11 | phosphoethanolamine transferase MCR | [5] |
| 6bnc | 18.4 | 3.4 | 257 | 335 | 11 | phosphoethanolamine transferase MCR | [5] |
| 6bne | 18.4 | 3.5 | 258 | 336 | 12 | phosphoethanolamine transferase MCR | [5] |
| 5i5f | 18.3 | 3.9 | 263 | 339 | 13 | inner membrane protein YejM | [4] |
| 6bnf | 18.3 | 3.4 | 256 | 336 | 12 | phosphoethanolamine transferase MCR | [5] |
| 5i5d | 18.3 | 3.8 | 263 | 339 | 13 | inner membrane protein YejM | [4] |
| 6fny | 18.2 | 3.8 | 268 | 509 | 13 | choline sulfatase BetC | [6] |
| 5mx9 | 18.1 | 3.4 | 257 | 324 | 13 | phosphatidylethanolamine transferase MCR-2 | [7] |
| 5gov | 18.0 | 3.4 | 256 | 321 | 12 | phosphatidylethanolamine transferase MCR-1 | [8] |
| 5k4p | 18.0 | 3.4 | 256 | 324 | 13 | phosphatidylethanolamine transferase MCR-1 | [9] |
| 5ylf | 17.9 | 3.5 | 257 | 330 | 12 | phosphatidylethanolamine transferase MCR-1 | [10] |
| 5grr | 17.9 | 3.5 | 256 | 323 | 13 | phosphatidylethanolamine transferase MCR-1 | [11] |
| 5lrn | 17.9 | 3.4 | 256 | 325 | 13 | phosphatidylethanolamine transferase MCR-1 | [12] |
| 1fsu | 17.8 | 3.9 | 273 | 475 | 13 | N-acetylgalactosamine-4-sulfatase | [13] |
| 5yle | 17.8 | 3.4 | 256 | 331 | 12 | phosphatidylethanolamine transferase MCR-1 | [10] |
| 5ylc | 17.8 | 3.4 | 256 | 330 | 12 | phosphatidylethanolamine transferase MCR-1 | [10] |

| | | | | | | | |
|---|---|---|---|---|---|---|---|
| 5zjv | 17.8 | 3.4 | 256 | 331 | 12 | phosphatidylethanolamine transferase MCR-1 | [14] |
| 5i5h | 17.7 | 3.7 | 249 | 327 | 12 | inner membrane protein YejM | [4] |
| 5lrm | 17.7 | 3.4 | 249 | 312 | 13 | phosphatidylethanolamine transferase MCR-1 | [12] |
| 4upk | 17.6 | 4.0 | 285 | 506 | 12 | phosphonate monoester hydrolase | NP |
| 3ed4 | 17.5 | 3.9 | 285 | 467 | 13 | arylsulfatase | NP |
| 4uph | 17.5 | 4.0 | 275 | 504 | 12 | phosphonate monoester hydrolase | NP |
| 6b0k | 17.5 | 4.0 | 268 | 452 | 13 | endo-4S-ι-carrageenan sulfatase | [15] |
| 4fdi | 17.4 | 3.8 | 274 | 494 | 13 | N-acetylgalactosamine-6-sulfatase | [16] |
| 4cyr | 17.4 | 4.0 | 279 | 526 | 12 | arylsulfatase | NP |
| 4fdj | 17.4 | 3.9 | 276 | 493 | 13 | N-acetylgalactosamine-6-sulfatase | [16] |
| 4tn0 | 17.4 | 3.3 | 244 | 308 | 13 | phosphoethanolamine transferase EptC | [17] |
| 6b1v | 17.4 | 4.0 | 274 | 452 | 12 | endo-4S-ι-carrageenan sulfatase | [15] |
| 2w8s | 17.2 | 4.2 | 281 | 513 | 11 | promiscuous hydrolase | [18] |
| 4upl | 17.2 | 4.0 | 290 | 553 | 11 | sulfatase | NP |
| 6b0j | 17.2 | 4.0 | 274 | 453 | 12 | endo-4S-ι-carrageenan sulfatase | [15] |
| 4miv | 17.1 | 3.7 | 265 | 484 | 12 | N-sulphoglucosamine sulphohydrolase | [19] |
| 6bia | 17.1 | 4.1 | 272 | 452 | 14 | endo-4S-ι-carrageenan sulfatase | [15] |
| 5fgn | 17.1 | 3.5 | 253 | 536 | 11 | lipooligosaccharide phosphoethanolamine transferase A | [20] |
| 4kay | 17.0 | 3.8 | 258 | 333 | 10 | lipooligosaccharide phosphoethanolamine transferase A | [21] |
| 4kav | 17.0 | 3.7 | 250 | 334 | 10 | lipooligosaccharide phosphoethanolamine transferase A | [21] |

[a]NP – no publication.

**Supplementary Table 2.** $^{1}$H and $^{13}$C NMR chemical shifts (ppm) at 50 °C of the GAC repeating unit and inter-residue correlations from $^{1}$H,$^{1}$H-NOESY and $^{1}$H,$^{13}$C-HMBC experiments.

| **Sugar residue** | | **$^{1}$H/$^{13}$C** | | | | | | | | **$^{31}$P** | **Correlation to atom (from anomeric atom)** | |
|---|---|---|---|---|---|---|---|---|---|---|---|---|
| | | 1 | 2 | 3 | 4 | 5 | 6 | Me | CO | $PO_4$ | NOE | HMBC |
| →2,3)-α-L-Rha*p*-(1→3) | **RI** | 5.10 [n.r.] | 4.29 | 4.00 | 3.54 | 3.83 | 1.33 | | | | H3, **RI** | C3, **RII** |
| | | 101.7{177} | 77.2 | 80.4 | 72.3 | 70.1 | 17.7 | | | | H5, **RII** | H3, **RII** |
| | **RI'** | 5.19 | 4.27 | | | | | | | | | |
| | | 101.5 | 78.3 | | | | | | | | | |
| →3)-α-L-Rha*p*-(1→2) | **RII** | 5.17 [n.r.] | 4.07 | 3.86 | 3.56 | 3.74 | 1.28 | | | | H2 **RI** | C2, **RI** |
| | | 102.2{178} | 70.7 | 77.9 | 72.6 | 70.2 | 17.4 | | | | | H2, **RI** |
| | **RII'** | 5.04 | 4.16 | | | | | | | | H2, **RI'** | C2, **RI'** |
| | | 102.8 | 70.8 | | | | | | | | | H2, **RI'** |
| β-D-Glc*p*NAc (1→3) | **GIII** | 4.76 [7.9] | 3.74 | 3.57 | 3.47 | 3.47 | 3.78, 3.94 | 2.05 | | | H3, **RI** | C3, **RI** |
| | | 103.2{167} | 56.8 | 74.8 | 70.8 | 76.6 | 61.8 | 23.2 | 175.3 | | | H3, **RI** |
| | **GIII'** | 4.77 | 3.78 | 3.59 | 3.48 | 3.59 | 4.10, 4.19 | | | | | |
| | | 103.0 | 56.7 | 74.7 | 70.6 | 75.4 | 65.4[b] | | | | | |
| D-Gro-1-*P* | **Gro** | 3.91, 3.96 | 3.93 | 3.64, 3.70 | | | | | | 1.2 | | |
| | | 67.3[a] | 71.6[b] | 63.1 | | | | | | | | |

$^{3}J_{HH}$ and $^{1}J_{CH}$ values are given in Hertz in square brackets and braces, respectively, where n.r. denotes not resolved; [a] $^{2}J_{P,\,C1}$ = 5.6 Hz, [b] $^{3}J_{P,\,C2}$ = 7.6 Hz.

**Supplementary Table 3.** Bacterial strains and plasmids.

| Strain or plasmid | Description | Reference |
|---|---|---|
| ***Bacteria*** | | |
| **GAS** | | |
| 5005 | GAS, M1T1-serotype strain | [22] |
| 5448 | GAS, M1T1-serotype strain | [23] |
| 2221 | GAS, M1T1-serotype strain | [22] |
| 5005Δ*gacH* | GAS 5005 has a 130-nt deletion in *gacH* | This study |
| 5005Δ*gacH*:*gacH* | 5005Δ*gacH* is complemented with WT *gacH* in *cis* | This study |
| 5005Δ*gacL* | GAS 5005 *gacL*:*aadA* deletion mutant (has a nonpolar *aadA* spectinomycin resistance cassette inserted in *gacL)* | [24] |
| 5005Δ*gacI* | GAS 5005 has a 82-nt deletion in *gacI* | [24] |
| 2221Δ*gacH* | GAS 2221 has a 130-nt deletion in *gacH* | This study |
| 2221Δ*gacH*:*gacH* | 2221Δ*gacH* is complemented with WT *gacH* in *cis* | This study |
| 5448Δ*gacI* | GAS 5448 *gacI:CAT* deletion mutant (has a nonpolar CAT chloramphenicol resistance cassette inserted in *gacH)* | [25] |
| 5448Δ*gacI:gacI* | 5448Δ*gacI* is complemented with WT *gacI* in *cis* | [25] |
| 5448Δ*gacH* | GAS 5448 *gacH:CAT* deletion mutant (has a nonpolar CAT chloramphenicol resistance cassette inserted in *gacH)* | This study |
| *5448ΔgacH*:p*gacH* | 5448Δ*gacH* is complemented with p*gacH*_erm carrying WT *gacH* | This study |
| 5448Δ*czcD* | GAS 5448 *czcD*:*aadA* deletion mutant | [26] |
| ***Streptococcus mutans*** | | |
| Xc | Serotype *c* strain | [27] |
| SMUΔ*sccH* | *S. mutans* Xc *sccH:erm* deletion mutant [has a nonpolar erythromycin (erm) resistance cassette inserted in *sccH*)] | This study |
| SMUΔ*sccH*:p*sccH* | SMUΔ*sccH* is complemented with p*sccH* carrying WT *sccH* | This study |
| SMUΔ*sccH*:p*gacH* | SMUΔ*sccH* is complemented with p*gacH*_cm carrying WT *gacH* | This study |
| ***Escherichia coli*** | | |
| *E. coli* DH5α | *E. coli* cells used for cloning | Invitrogen |
| *E. coli* Rosetta (DE3) | *E. coli* cells used for protein expression | Novagen |
| ***Plasmids*** | | |
| pBBL740 | GAS integrational plasmid | [28] |
| pBBL740Δ*gacH* | pBBL740 derived plasmid containing a 1.4 kbp DNA fragment carrying the *gacH* locus with a 130-nt deletion | This study |
| pBBL740*gacH* | pBBL740 derived plasmid containing a 1.4 kbp DNA fragment carrying the *gacH* locus | This study |
| pHY304 | A temperature sensitive *E.coli-Streptococcus* shuttle vector | [29] |
| pHY304Δ*gacH* | Derivative of pHY304 expressing CAT (a nonpolar CAT chloramphenicol resistance cassette) flanked with *gacH* 5' and 3' regions | This study |
| pET-NT | A modified pET-Duet1 (Novagen) vector that allows the creation of N-terminus His-tagged proteins with a TEV protease cleavage site | This study |
| pETGacH | pET-NT derived plasmid expressing extracellular domain of GacH fused with N-terminal TEV protease cleavable His-tag | This study |

| | | |
|---|---|---|
| pDC123 | *E. coli-streptococcus* shuttle vector, JS-3 replicon, CAT resistance cassette | [30] |
| pDCerm | pDC123 derivative with Erm (erm of Tn916ΔE) replacing CAT | [31] |
| p*sccH* | pDC123 derived plasmid expressing *sccH* | This study |
| p*gacH*_cm | pDC123 derived plasmid expressing *gacH* | This study |
| p*gacH*_erm | pDCerm derived plasmid expressing *gacH* | This study |

**Supplementary Table 4.** Primers.

| Primer | Sequence[a] |
|---|---|
| 5005-f | CGTCTGGATCCAATTGGTTAAATATTAGTCACC |
| gacHdel-f | GGACTTGCTATTTGTATTGTGGAAGGTAAAATATTATCTGGTC |
| gacHdel-r | GACCAGATAATATTTTACCTTCCACAATACAAATAGCAAGTCC |
| 5005-r | GCGCGCTCGAGGATACTTTTGAATCAGTATGCTC |
| gacH-NcoI-f | CGTGAGCCATGGAAAAACCGACAAATTATAGCC |
| gacH-XhoI-r | GCGCGCTCGAGTTAACGTGATATCTTAAAAAAAG |
| 5448-f | GCGCTCGAGGATCTTATTGTTTCACATATGTCGC |
| 5448CAT-r | GGTGGTATATCCAGTGATTTTTTTCTCCATTCATTTAAGTTTCTCCATTAAACGC |
| 5448CAT-f | TACTGCGATGAGTGGCAGGGCGGGGCGTAAGAAAAATGCTAGGCTGTCCG |
| 5448-r | GCGAAGCTTGAATGAATACCCAGGAAGTGAGAC |
| sccH-f | CTGCAAACAGTCAAGAGGCTCAG |
| sccH-erm-r | GTTTTGAGAATATTTTATATTTTTGTTCATGACTTTATTTTTTGCAGAGAACGCTT |
| sccH-erm-f | AGTTATCTATTATTTAACGGGAGGAAATAAATCTTTTTTTATGAAAAATACTTATGAGAA |
| sccH-r | GACGACTGTCAGAACCTGGAGAAA |
| sccH-c-f | CGCACATGTCAGAAATCAATTATCC |
| sccH-c-r | GAGACTAGCATTCTTGAAGCATATCC |
| gacH-EcoRI-f | CCGGAATTCATGATTAAAGACACATTTTTAAAAACCAAT |
| gacH-BglII-r | GAAGATCTTTAACGTGATATCTTAAAAAAAGTTTTGTGT |
| sccH-EcoRI-f | CCGGAATTCATGAAACAATTGAAAAAAATATGGGATA |
| sccH-BglII-r | GAAGATCTTTATCGAATGTCAAAGAATCCTTTATAGTT |
| oKrmit-Tnseq2 | CAAGCAGAAGACGGCATACGAAGCGCCTACGAGGAATTTGTATCG |
| AdapterPCR | AATGATACGGCGACCACCGAGATCACACTCTTTCCCTACACGACGCTCTTCC |

[a] - Restriction sites are underlined.

**Supplementary Table 5.** Data collection and refinement statistics.

| | GacH<br>PDB ID 5U9Z | GacH in complex *sn*-Gro-1-P<br>PDB ID 6DGM |
|---|---|---|
| **Data collection** | | |
| Wavelength (Å) | 0.9793 | 0.97946 |
| Space group | $P2_12_12_1$ | $P2_12_12_1$ |
| Cell dimensions: | | |
| *a*, *b*, *c* (Å) | 68.87, 78.17, 172.66 | 74.65 82.97 130.36 |
| α, β, γ (°) | 90, 90, 90 | 90, 90, 90 |
| Resolution (Å) | 49.5–2.00 (2.05–2.00)[a] | 39.5–1.49 (1.53–1.49) |
| $R_{sym}$ | 0.083 (0.827)[b] | 0.080 (1.143) |
| $CC_{1/2}$[c] | 99.3 (61.4) | 99.9 (49.6) |
| $I / \sigma I$ | 8.50 (1.11) | 14.23 (1.20) |
| Completeness (%) | 98.7 (90.0) | 96.3 (73.4) |
| Multiplicity | 3.6 (2.5) | 6.5 (5.0) |
| **Refinement** | | |
| Resolution (Å) | 49.5–2.00 | 39.5–1.49 |
| No. reflections (total / free) | 119860 / 5868 | 127526 / 6361 |
| $R_{work}$ / $R_{free}$ | 0.215 / 0.240 | 0.153 / 0.175 |
| Number of atoms: | | |
| Protein | 6058 | 6944 |
| Ligand/ion | 2 | 24 |
| Water | 321 | 846 |
| *B*-factors: | | |
| Protein | 50.2 | 21.3 |
| Ligand/ion | 36.3 | 26.9 |
| Water | 46.6 | 32.2 |
| All atoms | 50.3 | 22.7 |
| Wilson *B* | 40.3 | 24.3 |
| R.m.s. deviations: | | |
| Bond lengths (Å) | 0.003 | 0.008 |
| Bond angles (°) | 0.549 | 0.925 |
| Ramachandran distribution[d] (%): | | |
| Favored | 96.6 | 97.2 |
| Allowed | 3.4 | 2.8 |
| Outliers | 0 | 0 |
| Rotamer outliers[d] (%) | 0 | 0.44 |
| Clashscore[e] | 0.25 | 0.66 |
| MolProbity score[f] | 0.82 | 0.86 |

[a]Values in parentheses are for the highest-resolution shell.
[b]Friedel pairs are treated as different reflections.
[c]$CC_{1/2}$ correlation coefficient as defined in Karplus & Diederichs [32] and calculated by *XSCALE* [33].
[d]Calculated using the MolProbity server (http://molprobity.biochem.duke.edu) [34].
[e]Clashscore is the number of serious steric overlaps (> 0.4 Å) per 1000 atoms.
[f]MolProbity score combines the clashscore, rotamer, and Ramachandran evaluations into a single score, normalized to be on the same scale as X-ray resolution [34].

### Supplementary References


1. Lu, D. et al. Structure-based mechanism of lipoteichoic acid synthesis by *Staphylococcus aureus* LtaS. *Proc. Natl. Acad. Sci. USA* **106**, 1584-1589 (2009).
2. Schirner, K., Marles-Wright, J., Lewis, R.J. & Errington, J. Distinct and essential morphogenic functions for wall- and lipo-teichoic acids in *Bacillus subtilis*. *EMBO J.* **28**, 830-842 (2009).
3. Campeotto, I. et al. Structural and mechanistic insight into the *Listeria monocytogenes* two-enzyme lipoteichoic acid synthesis system. *J. Biol. Chem.* **289**, 28054-28069 (2014).
4. Dong, H. et al. Structural insights into cardiolipin transfer from the Inner membrane to the outer membrane by PbgA in Gram-negative bacteria. *Sci. Rep.* **6**, 30815 (2016).
5. Stogios, P.J. et al. Substrate recognition by a colistin resistance enzyme from *Moraxella catarrhalis*. *ACS Chem. Biol.* **13**, 1322-1332 (2018).
6. van Loo, B. et al. Structural and mechanistic analysis of the choline sulfatase from *Sinorhizobium melliloti*: A class I sulfatase specific for an alkyl sulfate ester. *J. Mol. Biol.* **430**, 1004-1023 (2018).
7. Coates, K., Walsh, T.R., Spencer, J. & Hinchliffe, P. 1.12 Å resolution crystal structure of the catalytic domain of the plasmid-mediated colistin resistance determinant MCR-2. *Acta Crystallogr. F Struct. Biol. Commun.* **73**, 443-449 (2017).
8. Hu, M. et al. Crystal structure of *Escherichia coli* originated MCR-1, a phosphoethanolamine transferase for colistin resistance. *Sci. Rep.* **6**, 38793 (2016).
9. Stojanoski, V. et al. Structure of the catalytic domain of the colistin resistance enzyme MCR-1. *BMC Biol.* **14**, 81 (2016).
10. Wei, P. et al. Substrate analog interaction with MCR-1 offers insight into the rising threat of the plasmid-mediated transferable colistin resistance. *FASEB J.* **32**, 1085-1098 (2018).
11. Ma, G., Zhu, Y., Yu, Z., Ahmad, A. & Zhang, H. High resolution crystal structure of the catalytic domain of MCR-1. *Sci. Rep.* **6**, 39540 (2016).
12. Hinchliffe, P. et al. Insights into the mechanistic basis of plasmid-mediated colistin resistance from crystal structures of the catalytic domain of MCR-1. *Sci. Rep.* **7**, 39392 (2017).
13. Bond, C.S. et al. Structure of a human lysosomal sulfatase. *Structure* **5**, 277-289 (1997).
14. Liu, Z.X., Han, Z., Yu, X.L., Wen, G. & Zeng, C. Crystal structure of the catalytic domain of MCR-1 (cMCR-1) in complex with D-xylose. *Crystals* **8**, 172 (2018).
15. Hettle, A.G. et al. The molecular basis of polysaccharide sulfatase activity and a nomenclature for catalytic subsites in this class of enzyme. *Structure* **26**, 747-758 (2018).
16. Rivera-Colon, Y., Schutsky, E.K., Kita, A.Z. & Garman, S.C. The structure of human GALNS reveals the molecular basis for mucopolysaccharidosis IV A. *J. Mol. Biol.* **423**, 736-751 (2012).
17. Fage, C.D., Brown, D.B., Boll, J.M., Keatinge-Clay, A.T. & Trent, M.S. Crystallographic study of the phosphoethanolamine transferase EptC required for polymyxin resistance and motility in *Campylobacter jejuni*. *Acta Crystallogr. D Biol. Crystallogr.* **70**, 2730-2739 (2014).
18. van Loo, B. et al. An efficient, multiply promiscuous hydrolase in the alkaline phosphatase superfamily. *Proc. Natl. Acad. Sci. USA* **107**, 2740-2745 (2010).
19. Sidhu, N.S. et al. Structure of sulfamidase provides insight into the molecular pathology of mucopolysaccharidosis IIIA. *Acta Crystallogr. D Biol. Crystallogr.* **70**, 1321-1335 (2014).
20. Anandan, A. et al. Structure of a lipid A phosphoethanolamine transferase suggests how conformational changes govern substrate binding. *Proc. Natl. Acad. Sci. USA* **114**, 2218-2223 (2017).
21. Wanty, C. et al. The structure of the neisserial lipooligosaccharide phosphoethanolamine transferase A (LptA) required for resistance to polymyxin. *J. Mol. Biol.* **425**, 3389-3402 (2013).
22. Sumby, P. et al. Evolutionary origin and emergence of a highly successful clone of serotype M1 group a Streptococcus involved multiple horizontal gene transfer events. *J. Infect. Dis.* **192**, 771-782 (2005).

23. Kansal, R.G., Nizet, V., Jeng, A., Chuang, W.J. & Kotb, M. Selective modulation of superantigen-induced responses by streptococcal cysteine protease. *J. Infect. Dis.* **187**, 398-407 (2003).
24. Rush, J.S. et al. The molecular mechanism of N-acetylglucosamine side-chain attachment to the Lancefield group A carbohydrate in *Streptococcus pyogenes*. *J. Biol. Chem.* **292**, 19441-19457 (2017).
25. van Sorge, N.M. et al. The classical Lancefield antigen of Group A Streptococcus is a virulence determinant with implications for vaccine design. *Cell Host Microbe* **15**, 729-740 (2014).
26. Ong, C.L., Gillen, C.M., Barnett, T.C., Walker, M.J. & McEwan, A.G. An antimicrobial role for zinc in innate immune defense against group A streptococcus. *J. Infect. Dis.* **209**, 1500-1508 (2014).
27. Koga, T., Asakawa, H., Okahashi, N. & Takahashi, I. Effect of subculturing on expression of a cell-surface protein antigen by *Streptococcus mutans*. *J. Gen. Microbiol.* **135**, 3199-3207 (1989).
28. Zhu, H., Liu, M., Sumby, P. & Lei, B. The secreted esterase of group a streptococcus is important for invasive skin infection and dissemination in mice. *Infect. Immun.* **77**, 5225-5232 (2009).
29. Chaffin, D.O., Beres, S.B., Yim, H.H. & Rubens, C.E. The serotype of type Ia and III group B streptococci is determined by the polymerase gene within the polycistronic capsule operon. *J. Bacteriol.* **182**, 4466-4477 (2000).
30. Chaffin, D.O. & Rubens, C.E. Blue/white screening of recombinant plasmids in Gram-positive bacteria by interruption of alkaline phosphatase gene (*phoZ*) expression. *Gene* **219**, 91-99 (1998).
31. Jeng, A. et al. Molecular genetic analysis of a group A Streptococcus operon encoding serum opacity factor and a novel fibronectin-binding protein, SfbX. *J. Bacteriol.* **185**, 1208-1217 (2003).
32. Karplus, P.A. & Diederichs, K. Linking crystallographic model and data quality. *Science* **336**, 1030-1033 (2012).
33. Kabsch, W. *Xds*. *Acta Crystallogr. D Biol. Crystallogr.* **66**, 125-132 (2010).
34. Chen, V.B. et al. *MolProbity*: all-atom structure validation for macromolecular crystallography. *Acta Crystallogr. D Biol. Crystallogr.* **66**, 12-21 (2010).
